# Parental haplotypes reconstruction in up to 440,209 individuals reveals recent assortative mating dynamics

**DOI:** 10.1101/2025.09.24.678243

**Authors:** Robin J. Hofmeister, Davide Marnetto, Théo Cavinato, Leona Knüsel, Giorgia Modenini, Erwan Pennarun, Estonian Biobank research team, Nicola Barban, Reedik Mägi, Tabea Schoeler, Luca Pagani, Zoltán Kutalik

## Abstract

Assortative mating (AM), the tendency for individuals to choose partners with similar traits, plays an important role in social stratification and has a wide-spread impact on the genetic architecture of complex human traits. A genetic footprint of this behaviour, genetic assortative mating (GAM), has been documented for many traits. However, most existing approaches to estimate GAM either rely on genotyped couples – rare in large biobanks – or on estimates based on gametic phase disequilibrium (GPD), which reflect cumulative effects over multiple generations and cannot detect short-term events.

We introduce a novel, scalable approach that infers genetic data for the parental generation to boost statistical power and improve interpretability when estimating GAM in biobank cohorts. Our method reconstructs the two parental haplotypes of biobank individuals by leveraging state-of-the-art inter-chromosomal phasing based on close relative information. By correlating polygenic scores computed separately on maternally and paternally inherited haplotypes, we infer the extent of GAM.

Applied to 245,884 individuals from the UK Biobank and 194,325 individuals from the Estonian Biobank, our haplotype-based estimates showed strong concordance with GAM estimates from genotyped mate pairs across 69 traits and improved performance compared to existing GPD-based methods. We replicated previously identified patterns of GAM for many traits (educational attainment, height, BMI, and alcohol consumption), and revealed new ones (overall health and sedentary lifestyle). Temporal and geographic stratification revealed accelerated assortment in recent generations — particularly for education and height — and modest differences between urban and rural contexts.

Our method enables scalable, interpretable, and generation-specific estimation of GAM in large biobank cohorts, providing new insights into the dynamic nature of mate choice and its impact on the genetic makeup of populations.

## Introduction

Assortative mating (AM) is the tendency for individuals to select partners with similar characteristics. It plays an important role in shaping human behaviour, social structure and genetic architecture of human populations^1,2,3^. It has been widely documented in humans for a broad range of phenotypes, including height, body mass index (BMI), educational attainment, income, lifestyle behaviors, and even disease risk^4,5,6,7,8^.

This phenotypic similarity between partners has important implications across disciplines. In social sciences, AM and homogamy have long been studied as a mechanism contributing to the closure of social groups and the emergence of growing inequality over generations, and fragmentation in modern societies^9,10^. Conversely, intermarriage is seen as a driver of cultural integration, reducing distinctions between groups over generations^11^.

While AM is often assumed to be driven by the active choice of a partner with similar phenotypes, it can emerge through different alternative mechanisms: Individuals tend to form couples with those who come from similar social backgrounds (social homogamy^12^) or with those they encounter more frequently in daily environments (contextual/structural opportunity^13^). For example, temporal and spatial constraints^14^ can greatly limit the pool of potential partners, producing apparent similarities that reflect local (e.g., neighborhood composition, urban versus rural settings) and temporal (e.g., life stage, prevailing cultural norms or economic conditions) environments rather than active choice. Moreover, indirect assortment for one trait can emerge when couples directly assort for a correlated trait^15,16,17^. Finally, cohabiting partners may become more similar through mutual influence, although such effects are difficult to detect empirically^17,18^. The various sources leading to AM can complicate the interpretation of the observed patterns, and separating them is a nontrivial task^17^.

AM has well-characterised consequences for genetics. Genetic variants associated with traits under assortment will themselves be subject to indirect assortment, leading to genetic similarity between mate pairs. As a consequence, AM inflates not only phenotypic, but also genetic variance in the population, strengthens correlations among relatives, and leads to deviations from Hardy–Weinberg equilibrium (HWE) at causal loci^19,20^. As a consequence, AM leads to an increased prevalence of autosomal recessive diseases^21^. It also generates correlations between trait-associated alleles (TAAs) at unlinked loci, termed gametic phase disequilibrium (GPD), which differs from local linkage disequilibrium, as it reflects long-range, directional correlations accumulated over generations^7^. GPD can bias genetic association studies by violating the assumption of independence among unlinked loci. This can lead to inflated GWAS test statistics, biased SNP effect estimates, and overestimated SNP-based heritability and genetic correlations^22,23^. Such biases have implications for downstream analyses, such as Mendelian Randomisation, especially for trait pairs strongly affected by AM. Estimating AM in humans is therefore essential not only for inferring evolutionary events and patterns of human behaviour, but also for evaluating risk of bias in genetic epidemiology investigations.

Several recent studies have attempted to quantify genetic assortative mating (GAM) by measuring genomic similarity between partners in several datasets using genome-wide SNP data^6,8^. While informative, these approaches require large numbers of genotyped couples, rarely available in biobanks.

An alternative strategy to estimate GAM without spousal data is to test for excess homozygosity at trait-associated loci in unrelated individuals^24^. However, for highly polygenic traits, the loci driving assortment are likely numerous and widely dispersed across the genome, often on different chromosomes. As a result, the expected increase in homozygosity in the offspring is minimal, making this approach underpowered to detect GAM for polygenic traits. To address these limitations, Yengo et al.^7^ proposed estimating AM by quantifying GPD between TAAs in large population samples. This approach bypasses the need for couple genotypes and enables genome-wide assessment of AM at scale. In addition, detecting GPD is more tractable than testing for excess homozygosity, as it is designed to aggregate signals across a much larger number of alleles. However, because GPD assumes stability of both the genetic architecture of the studied trait and its AM pattern over generations, it builds up over time. Hence, this method captures only long-term historical assortment patterns. Therefore, estimating the genetic consequences of recent AM without genotype data of mate-pairs remains a major methodological challenge.

Here, we propose a novel approach to estimate recent GAM at biobank scale without requiring mate-pair information. Leveraging recent advances in inter-chromosomal phasing^25^, we reconstructed the maternally and paternally inherited haplotypes of up to a total of 440,209 individuals in the UK Biobank (UKBB) and Estonian Biobank (EstBB). We then computed polygenic scores (PGS) separately on each parental haplotype, effectively capturing the transmitted part of parental genomes. By correlating paternal and maternal haplotype-based PGS across individuals — which serve as imperfect proxies for parental pairs — we estimated the degree of assortative mating that occurred directly in the parental generation. Our analysis opens new avenues for estimating AM patterns for any heritable trait without couples and phenotypes and enabled testing AM differences across generations and populations.

## Results

### Method overview and performance

Assortative mating leaves detectable imprints in the offspring genome: when parents are genetically similar for a trait, the alleles they transmit tend to align more than expected under random mating, resulting in similar distributions of trait-associated alleles on the maternal and paternal haplotypes. Consequently, PGS computed from the two parental haplotypes of an individual are expected to correlate. Exploiting this phenomenon, we can estimate GAM by correlating maternal and paternal haplotype-based PGS across individuals.

To implement this approach, we first reconstructed and separated maternally and paternally inherited haplotypes in up to 245,884 white British individuals from UKBB and 194,325 individuals from the EstBB (Figure 1). This was achieved using recent advances in inter-chromosomal phasing, which leverages identity-by-descent (IBD) sharing with close relatives^25,26^.

**Fig. 1.**
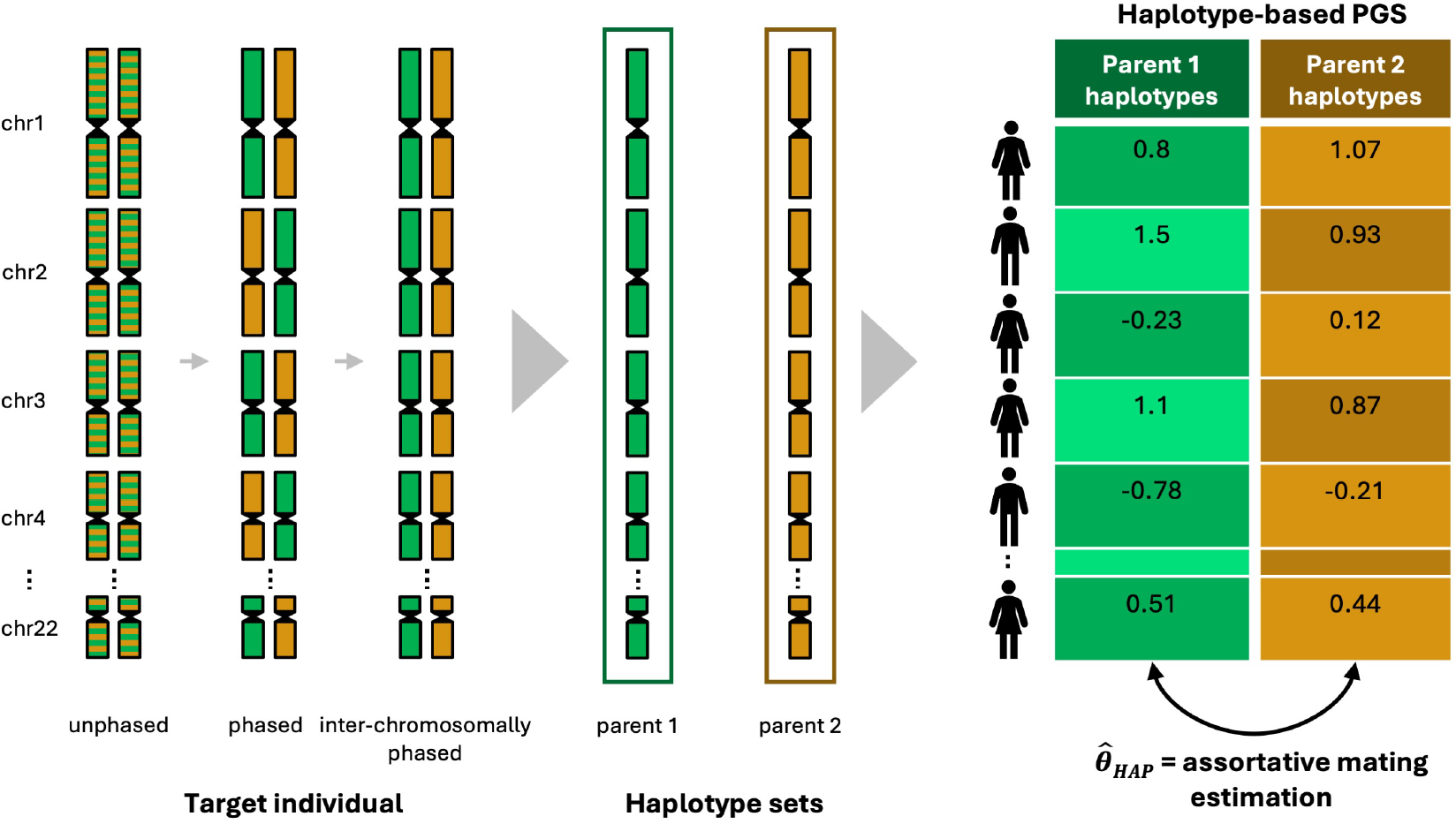
Haplotype-based Genetic Assortative Mating (GAM) estimation method. Schematic overview of the haplotype-based GAM estimation approach. For each target individual, unphased genotype data (left) are phased and then inter-chromosomally phased to separate the maternally and paternally inherited haplotypes. These two haplotype sets represent the genetic contributions from the two parents. PGS are computed separately for each parental haplotype set across all individuals (right), yielding two PGS values per individual, one per parent. The correlation between partial parental PGS values across the sample, 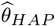, provides an estimate of assortative mating in the parental generation.

We then computed out-of-sample polygenic risk scores for a curated set of 69 complex traits (Supplementary Figure 1, see Methods), separately for maternally and paternally inherited haplotypes. Finally, we regressed the maternal haplotype-based PGS on the paternal haplotype-based PGS across individuals to estimate GAM in the parental generation (Figure 1, see Methods).

To assess the performance of our approach, we compared our GAM estimates across 69 traits to those derived from genotyped mate pairs using 22,068 couples in the UKBB and 10,618 couples in the EstBB as ground truth. We demonstrated via theoretical derivation and empirical validation that the assortment strength estimated by our method is half of the value that could be obtained when only using those samples for which parental genomes are available (*y* = 0.49*x* + 0; see Supplementary Notes 1 and Supplementary Figure 2). Furthermore, our approach showed strong agreement with these reference estimates in both biobanks (Supplementary Figure 3), achieving better trait ranking than the GPD-based approach (*r*^2^ = 0.737 and 0.638 in the UKBB and EstBB, respectively, *vs*. 0.731 and 0.479 for the GPD-based approach), lower estimation errors (RMSE= 0.0136 and 0.0132 *vs*. 0.0142 and 0.0150), and better calibration, with regression slopes closer to the expected line of slope 0.5 (*y* = 0.32*x* + 0 and *y* = 0.30*x* + 0 *vs. y* = 0.28*x* + 0 and *y* = 0.19*x* + 0), while preserving the same true positive rate (TPR) as the GPD-based method (TPR = 0.8 and 1 for both methods in the UKB and EstBB, respectively), defined as the proportion of traits under significant GAM (*P* − *value <* 0.05*/*60, see Methods) in the mate-pair data that were recovered by the haplotype-based or GPD-based approach.

To evaluate the influence of traits with strong GAM, we repeated the analysis after excluding height- and education-related traits (standing height, education, forced vital capacity FVC, fluid intelligence score, FEV1, Body mass index). The correlations with mate-pair estimates remained higher for our method, with *r*^2^ = 0.38 *vs*. 0.36 in the UKBB and 0.21 *vs*. 0.034 in the EstBB.

### GAM in the UK and Estonian populations

We characterised the landscape of GAM across 69 complex traits in the UK and Estonian populations using our haplotype-based method, and compared these estimates to those obtained from mate-pairs and from the GPD-based approach. Overall, GAM patterns were remarkably consistent between the two cohorts (*r* = 0.85, Figure 2A), indicating broadly similar trait priorities between the two countries when it comes to mate choice. Although less concordant, the same analysis with the GPD-based approach confirmed this trend (*r* = 0.67, Supplementary Figure 4).

**Fig. 2.**
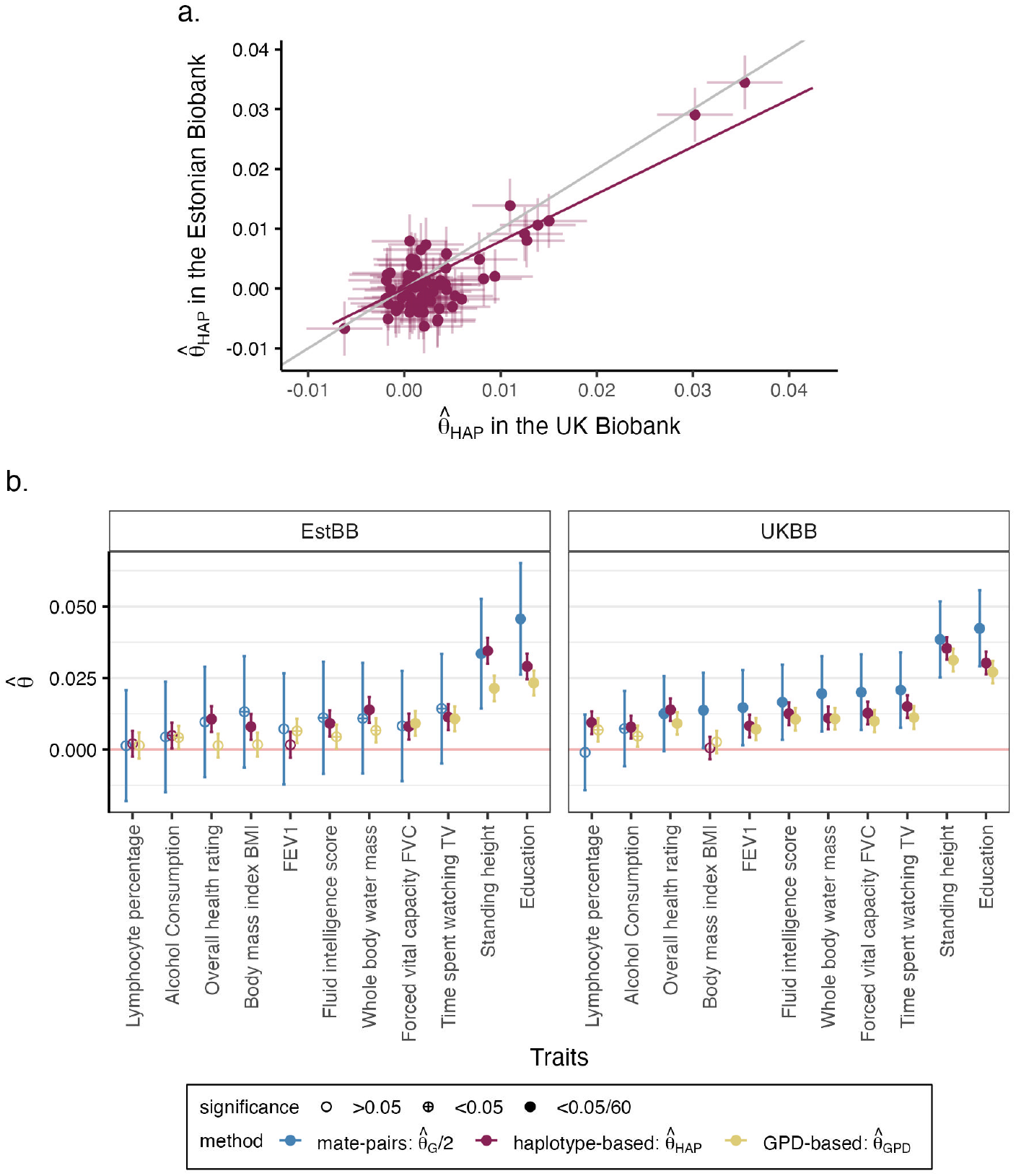
GAM estimates in the UK and Estonian cohorts. **a)** GAM computed with our haplotype-based approach 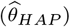 in the UK (x-axis) *vs*. Estonian Biobank (y-axis). Each dot show the 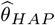 for a different trait, with error bars indicating 95% confidence intervals computed as 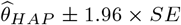. Grey line shows the identity line. Purple line shows the regression line *y* = 0.86*x* + 0. **b)** GAM estimates (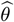, y-axis; blue for mate-pair 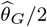 (rescaled for comparability); purple for haplotype-based 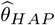; yellow for GPD-based 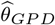) across 11 traits (x-axis) showing significant 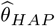 in at least one cohort. Shapes indicate non-significant (empty circle), nominally significant (crossed circle) and Bonferroni significant (full circle) traits. Error bars show 95% confidence intervals computed as 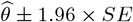.

We identified a total of 11 traits under significant GAM (*P-value* < 0.05/60, see Methods) using our approach (Supplementary Table 1: 10 in the UK- and 8 in the Estonian cohort, respectively (Figure 2B). Seven traits showed significant GAM in both cohorts: educational attainment, standing height, time spent watching TV, fluid intelligence score, whole body water mass, overall health rating, and forced vital capacity (FVC). These largely reflect established patterns of assortment across cognitive, behavioral, and anthropometric domains. Notably, two of these traits — fluid intelligence score and overall health rating — were not detected as significant by the GPD-based method in the EstBB (Supplementary Figure 5), underscoring the increased sensitivity of our haplotype-based approach.

We also observed cohort-specific signals: partners significantly assorted for BMI only in the Estonian cohort, detected only by our method. Conversely, three traits were significant only in the UK cohort: forced expiratory volume (FEV1), alcohol consumption, and lymphocyte percentage, with the latter two having been uniquely identified by our method (Figure 2B, Supplementary Figure 5).

Results from mate-pairs analysis showed partial overlap with our findings: in the UK Biobank, it identified nine significant traits, including one our method did not capture (BMI), but failed to detect assortment for alcohol consumption; in the Estonian Biobank, it identified only two significant traits, standing height and education attainment.

In contrast, the GPD-based approach identified two additional traits under significant GAM: hair colour in both cohorts – but with opposite directions of assortment – and age at first sexual intercourse in the UK Biobank. However, for hair colour, the estimates from the other methods were non-significant, yet consistently negative, suggesting that the GPD-based signal may reflect noise or long-term population stratification rather than recent assortative mating (Supplementary Figure 5).

To explore whether some of the observed assortment patterns are due to indirect assortment emerging through these traits being correlation with height or education (showing the strongest signals for assortative mating). To test this, we re-estimated GAM after conditioning the focal traits on these two traits (Supplementary Figure 6). In the UK Biobank, three traits remained significant using our approach: time spent watching TV, fluid intelligence score, and overall health rating. In contrast, only time spent watching TV was significant in the mate-pairs analysis, and all lost significance using the GPD-based approach. In the Estonian cohort, none remained significant regardless of the applied method.

Finally, to assess whether partner similarity may partly reflect shared environment, we additionally paired individuals based solely on geographic proximity (geo-pairs) and re-estimated PGS correlations. These geo-pairs showed weaker resemblance than real couples (Supplementary Figure 7), suggesting that while shared geography contributes to partner similarity, it does not fully account for the strength of assortative mating.

### Changes in assortative mating across generations

To investigate how assortative mating has changed over time, we leveraged a key advantage of our haplotype-based approach: it estimates assortment specifically in the parental generation of sampled individuals, rather than averaging patterns across multiple generations (like GPD-based approaches). By stratifying participants into year-of-birth (YoB) bins (Supplementary Figure 8), we were able to capture generational shifts in mate choice patterns with finer temporal resolution.

In both the UK and Estonian populations, we observed a general trend of intensified assortative mating in more recent generations: all traits with significant GAM in the non-stratified analysis showed increasing GAM with later YOB (10/10 in UKBB and 8/8 in EstBB; Figure 3). A similar trend was observed across all traits tested in the UKBB (60/69), but not overwhelmingly so in the EstBB (39/69, Supplementary Figure 9).

**Fig. 3.**
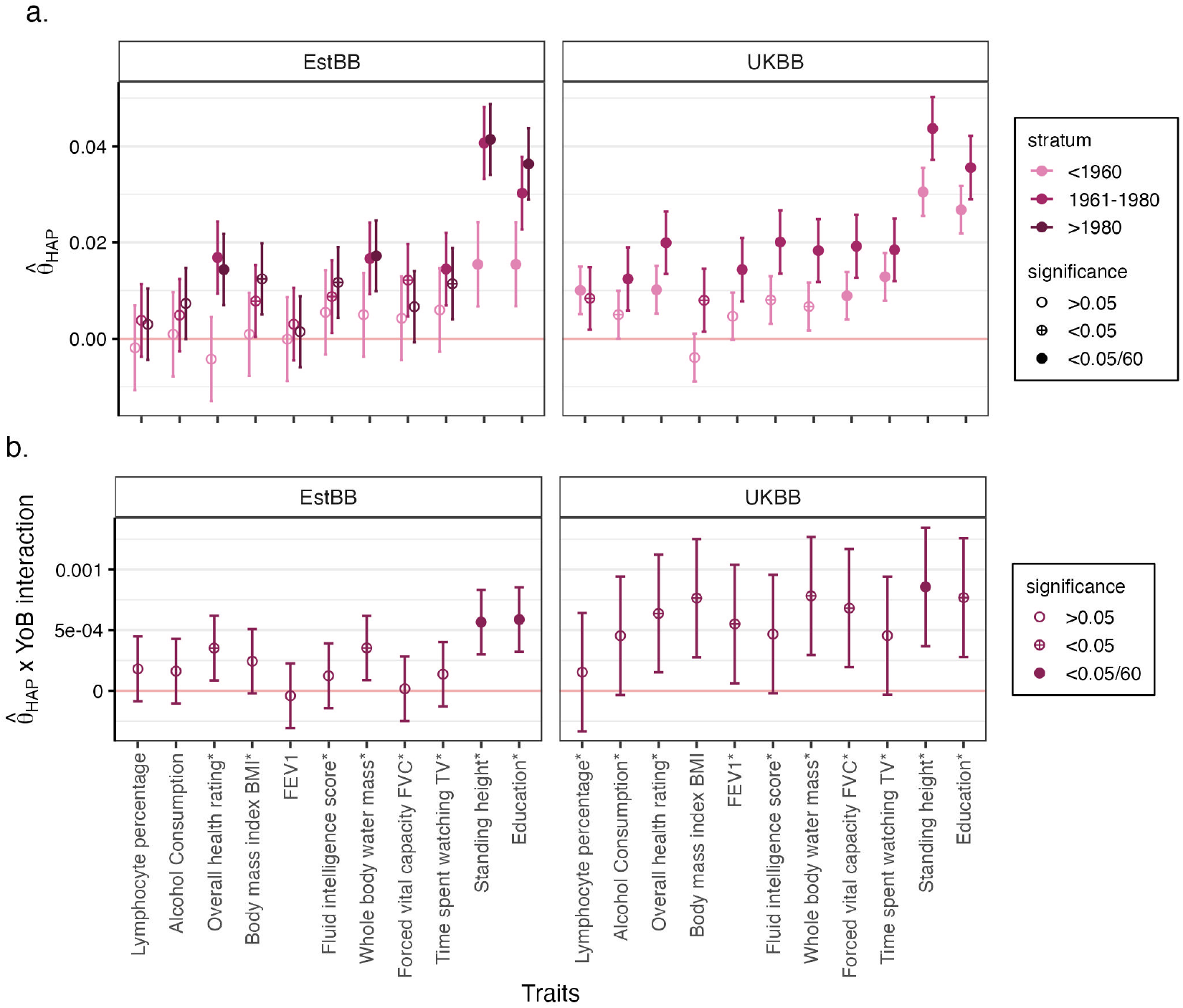
GAM estimates per Year of Birth (YoB) bins. **a)** Haplotype-based GAM (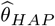, y-axis) per YoB bins in the UK and Estonian Biobank across the 11 traits showing significant 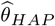 in at least one cohort (x-axis). Dot colors represent YoB bins, with darker colors indicating more recent generations. **b)** YoB interaction effect on 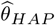 (y-axis) across traits (x-axis). For a) and b), shapes indicate non-significant (empty circle), nominally significant (crossed circle) and Bonferroni significant (full circle) traits. Error bars show 95% confidence intervals computed as 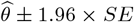. * indicate traits showing significant 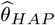 in the corresponding cohort (Figure 2).

Height showed a significant positive interaction with YoB in both cohorts, supporting a linearly increasing trend in assortment over time (Figure 3). Educational attainment also showed a significant generational increase in the EstBB, although no such pattern was detected in the UKBB.

Beyond these, many traits exhibited nominally significant interactions with YoB, suggesting a more subtle but widespread amplification of assortative behaviour (Supplementary Figure 9 and Supplementary Figure 10). In particular, in the Estonian population, hip circumference and smoking status showed evidence of changing assortment, but not linearly with YoB (Supplementary Figure 9): GAM estimates were significant only in the youngest cohort (YoB *>* 1980, Supplementary Figure 10), suggesting a recent acceleration in the assortment for these traits. A similar trend for hip circumference, but not for smoking status, was also observed in the UKBB. In contrast, the UK population exhibited an intensified GAM for alcohol consumption, with nominally significant linear YoB effect (Supplementary Figure 9) and stronger AM in the most recent generation (Figure 3).

For comparison, the GPD-based approach yielded broadly consistent results, but reflected long-term average trends over multiple generations rather than generation-specific changes revealed by our method (Supplementary Figure 9 and Supplementary Figure 11). Consequently, it detected only the significant interaction with YoB for standing height in the EstBB, failing to identify the other significant interactions revealed by the haplotype-based approach (*p >* 0.05). However, it did detect a significant interaction with YoB for whole body water mass (*p* = 5.12 *×* 10^−4^), for which our method showed only nominal significance (*p* = 8.78 *×* 10^−3^). This discrepancy likely reflects that the change occurred in older generations than the particpants’ parents, which cannot be captured by our haplotype-based approach, since it does not average GAM over multiple generations.

### Geographic variation in assortative mating

Next, we explored whether GAM varies by the settlement type of an individual’s residence in the UK and Estonian populations. Using the participants’ current address, we grouped them into categories ranging from rural (isolated dwelling, villages) to urban (town, city) environments. We then estimated GAM within each group and also tested for GAM trends by degree of urbanisation.

Overall, geographic effects on assortative mating were modest (Figure 4, Supplementary Figures 13 and 12). Although no geography effects reached Bonferroni-corrected significance, some traits exhibited notable patterns across geographic categories (Supplementary Figure 13): assortment on standing height was stronger in more rural areas, while educational attainment tended to show stronger assortment in urban areas. In the UK cohort, serum urate and calcium levels also showed higher GAM in rural areas, which may reflect indirect assortment through correlated traits, such as diet.

**Fig. 4.**
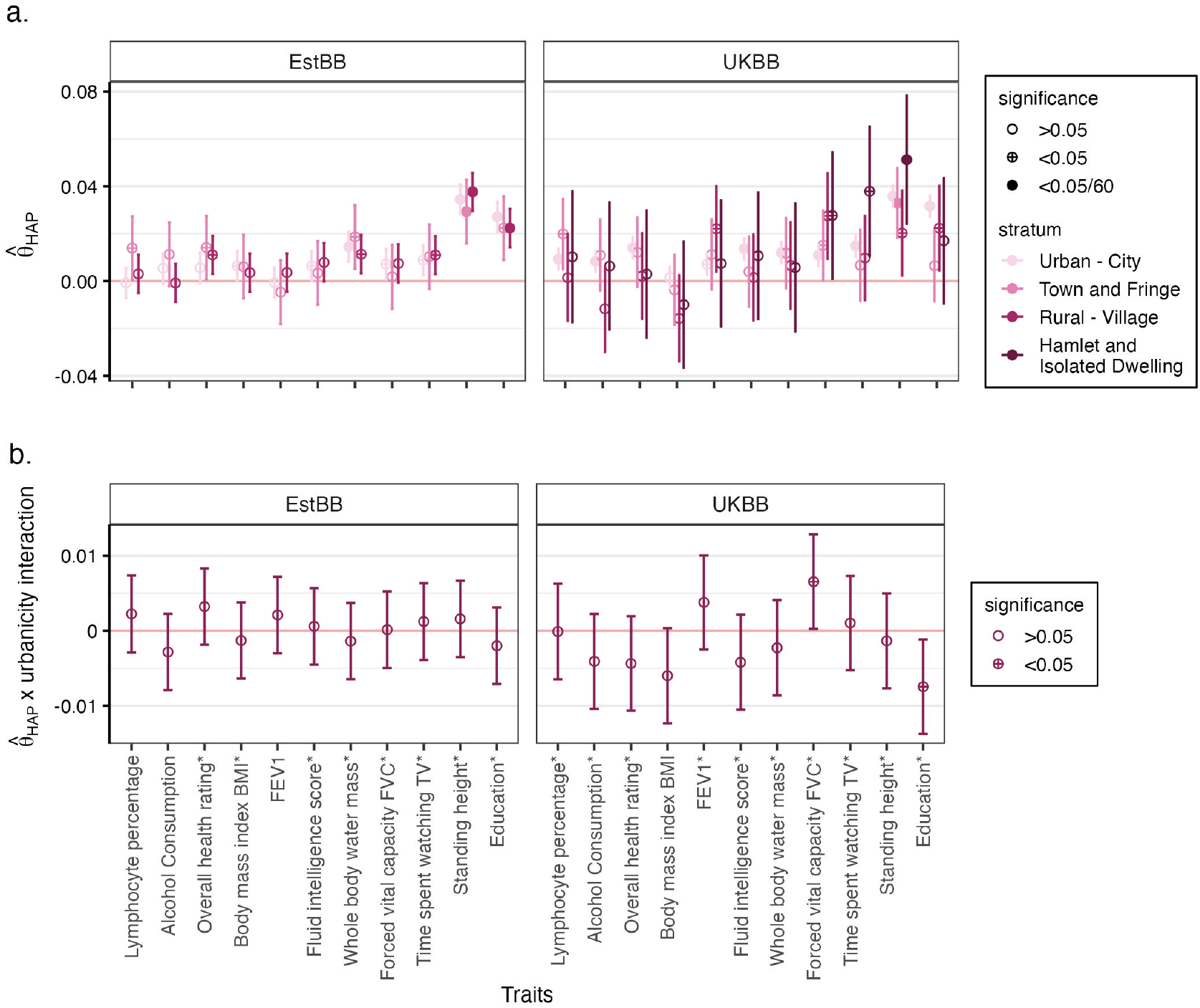
GAM estimates per settlement type. **a)** Haplotype-based GAM (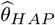, y-axis) per settlement type in the UK and Estonian Biobank across the 11 traits showing significant 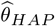 in at least one cohort (x-axis). Dot colors represent settlement type, with darker colors indicating more rural areas. **b)** Urbanicity interaction effect on 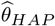 (y-axis) across traits (x-axis). For **a)** and **b)**, shapes indicate non-significant (empty circle), nominally significant (crossed circle) and Bonferroni significant (full circle) traits. Error bars show 95% confidence intervals computed as 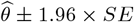. * indicate traits showing significant 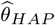 in the corresponding cohort (Figure 2).

## Discussion

We introduced a novel approach to estimate genetic assortative mating (GAM) at biobank scale by leveraging state-of-the-art inter-chromosomal phasing to reconstruct parental haplotypes. By comparing an individual’s PGS computed from maternal vs paternal haplotypes, our approach provides direct estimates of GAM between the participant’s parents without requiring their genotypes to be measured. Applied to the UK and Estonian Biobanks, we reconstructed parental haplotypes for roughly 50% of the UK Biobank (245,884 out of 500,000 individuals) and for almost the entire Estonian Biobank (194,325 out of 210,000 individuals). This enabled robust estimation of assortative mating for a specific generation across a broad range of traits, allowing for capturing temporal and geographic variations in mate choice.

Our method advances beyond existing approaches by focusing on single-generation assortment patterns, rather than the long-term average effects captured by GPD-based methods. This generational resolution facilitates the identification of punctual changes in AM patterns. Compared to couple-based analyses, our approach is scalable to hundreds of thousands of individuals and is neither constrained by the reliability of couple information nor the availability of their genotype data. Notably, our approach aligns more closely with mate-pair estimates than GPD-based methods, possibly because it reflects the most recent parental generation rather than long-term historical averages. This distinction is crucial when studying traits subject to rapid societal or environmental changes. As a sensitivity check, we also calculated similarity for “geo-pairs” (pairs of individuals living less than 1km from each other), revealing similarity induced by shared geographic environment (Supplementary Figure 7). This illustrates that partners are more similar than people living next door.

Our approach also has several limitations. First, it requires a certain degree of relatedness in the data to enable accurate inter-chromosomal phasing, which may limit its applicability in population cohorts with limited numbers of close relatives. Second, it captures assortment only among couples who had at least one child, and therefore does not fully reflect mating patterns in the entire population. Third, like the GPD-based approach, our method depends on the predictive performance of PGS, restricting its use to traits with sufficiently predictive scores and reducing power in populations where PGS portability is lower. For this reason, we applied the method only to non-admixed individuals, where the same scoring files could be used to calculate PGS on both haplotypes with comparable predictive performance. In admixed individuals, the two haplotypes may originate from different ancestral populations with different predictive accuracy, complicating interpretation. Future extensions of the approach to incorporate ancestry-aware PGS could enable GAM analyses in admixed populations and reveal novel, cross-population-specific mate preferences. Along these lines, GAM estimates may be influenced by population stratification despite the fact that we controlled for ancestry principal components when correlating the polygenic scores. Fourth, it is important to note that GAM does not necessarily translate into assortative mating at the phenotypic level: BMI, for example, shows stronger environmental assortment, whereas educational attainment and fluid intelligence exhibit stronger genetic assortment, likely reflecting the difference between genetically and environmentally correlated traits. Finally, the method does not distinguish between direct, indirect or confounded assortment, and thus some signals may reflect assortment on genetically correlated traits rather than the focal trait itself.

Our results confirm well-established patterns of assortative mating for traits such as educational attainment, standing height, and BMI, and also corroborate more recently reported signals, such as alcohol consumption^27,28^. Overall, the close agreement between our GAM estimates in the UK and Estonian populations — and with previous reports — suggests that mate choice for traits such as BMI, height, and educational attainment may be a widespread pattern in European populations, and possibly beyond. Understanding the social and historical factors underlying this phenomenon could provide valuable insight into its origins, while also highlighting the need for caution in GWAS for traits subject to assortative mating as they may generate spurious associations. In addition, we detect novel traits showing evidence of GAM, such as lymphocyte count, which however may reflect indirect assortment secondary to smoking^29^.

Crucially, we found that assortative mating has intensified in more recent cohorts^30,31^, particularly for height and education, consistent with relaxation of social constraints, changing social preferences and demographic trends over the 20^*th*^ century. As a result, bias in genetic analyses arising from assortative mating may be amplified in emerging biobank studies involving younger generations.

Using the same framework, we also examined geographic variation in GAM, although we acknowledge that UK and Estonian urbanisation patterns may differ substantially. While no strong regional differences emerged overall, we detected opposing patterns for education and height, with stronger assortment for education in urban areas and the opposite seems to hold for height.

Finally, while our GAM estimates were consistent between the UK and Estonian populations, the magnitudes of PGS correlations are not directly comparable due to differences in PGS predictive power and cohort composition. The better concordance with couple-based estimates in the EstBB may result from more reliable identification of couples based on trios, whereas UKBB couples were inferred indirectly from less precise criteria (see Methods).

In summary, our haplotype-based approach provides a scalable, interpretable, and generation-specific method to estimate GAM, revealing subtle yet meaningful differences in assortative mating across time, space, and traits. These findings underscore the complex interplay between social behavior and genetic architecture, opening new avenues for research into the evolutionary and societal drivers of mate choice, which appear to be widespread across multiple European cohorts and perhaps even to other cohorts worldwide. Importantly, the consequences of assortative mating on genetic architecture should not be overlooked. Although assortment is typically measured for common variants, as captured by PGS, its impact also propagates to rare variants. This can even induce correlations between common variant predisposition and rare variant risk in neurodevelopmental conditions^32^. As biobanks continue to expand and include more close relatives, we will be able to increase sample size enabling the detection of more subtle changes in assortative behaviour.

## Methods

### UK and Estonian Biobanks genotype data processing

We quality-controlled, phased, inter-chromosomally phased, and imputed the data as previously described for both biobanks^25^. This resulted in 245,884 white British UK Biobank individuals and 194,325 Estonian Biobank individuals that entered in our analyses.

### Mate-pairs identification

In the UK Biobank, we identified mate-pairs as suggested by Yengo *et al*.^7^. Briefly, we matched individuals into pairs based on: home location east and north coordinates, townsend deprivation index, number of smokers in household, number in household, length of time at the current address, average total household income before tax, living with husband or wife or partner, opposite sex, white British ancestry (data field #22006) and being unrelated. This resulted in 22,068 pairs.

In the Estonian Biobank, we used pairwise kinship estimates computed using the KING software v2.2.4^33^ together with age and sex to identify parent-offspring trios as having: kinship between 0.1767 and 0.3535, IBS0 below 0.0012, age difference greater than 15 years, and different sex of the parents^25^. It resulted in 14,063 trios for 10,618 unique pairs of parents.

### Polygenic score calculations and phenotype selection

We used a set of 146,389 white British individuals from the UK Biobank (data field #22006), distinct from the 245,884 individuals on whom we performed inter-chromosomal phasing, to derive polygenic weights as detailed below.

We selected a set of 81 traits and processed them as previously described^25^: for each individual, we averaged phenotype measurements across all available time points to obtain a single value per trait. Each trait was then inverse-normal quantile transformed using the `rntransform` function from the GENABEL R package^34^. For type 2 diabetes, cases were defined by the ICD-10 code `E11` (non-insulin-dependent diabetes mellitus), excluding individuals with the `E10` code (insulin-dependent diabetes mellitus) from both cases and controls to enhance specificity. Individuals diagnosed with other forms of diabetes (e.g., gestational diabetes) were excluded from both groups.

We conducted association testing for each trait using Regenie v3.2.9^35^. As recommended, only directly genotyped variants were included for model fitting (*step 1*), while imputed genotype dosages (field #22828, version 3) were tested in the association model (*step 2*).

We used LDpred2 (bigsnpr v.1.12.18)^36,37^ to adjust the summary statistics based on LD and to produce scoring files. As recommended in the tutorial of the tool, we restricted our analysis to the HapMap3+ variants and used the corresponding (precomputed) LD matrices covering these SNPs. First, we used the `snp_ldsc` function to estimate heritability. Then, we provided these estimates to the `snp_ldpred2_auto` function with the following parameters: burn_in=500, num_iter=500, report_step=20, allow_jump_sign=F, use_MLE=T, shrink_corrr=0.95, vec_p_init = seq_log(1e-4, 0.2, length.out = 10). We next use `pgs_calc` v1.6.0^38^ with default parameters to compute PGS from the previously produced scoring files.

We estimated the phenotypic variance explained by the PGS (*r*^2^) in an out-of-sample set of 245,884 white British UK Biobank individuals (i.e., those not used for association testing), and kept only the 69 traits with *r*^2^ *>* 0.01 for downstream analysis (Supplementary Figure 1).

To account for the number of traits tested in downstream analyses, we computed phenotypic correlation between each pair of traits, and used the function *simpleM* from the R package *hscovar* to compute the effective number of traits (*N*_*eff*_ (*traits*) = 60), which was then used for the Bonferroni correction.

### Haplotype-based PGS calculation

To estimate haplotype-based PGS for our set of inter-chromosomally phased individuals, we encoded parental haplotype in separate files. Let us consider an imputed variant *v* for a target individual *t* with haploid imputed dosages 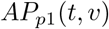 and 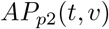 for parent 1 and parent 2 haplotype, respectively. The genotype probability triplets (*p*_*AA*_, *p*_*AB*_, *p*_*BB*_) were then defined as follows:

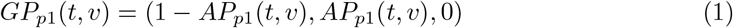

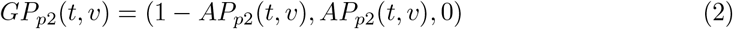

### Mate pair-based assortative mating calculations

We estimated phenotypic assortative mating between mate-pairs by regressing the trait of the first individual *T*_*m*1_ on the trait of the second individual *T*_*m*2_:

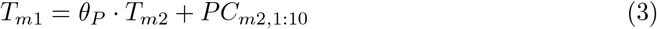

where *θ*_*P*_ represents the phenotypic AM, and *PC*_*m*2,1:10_ are the first 10 ancestry principal components of the partner individual included to account for population structure.

Similarly, we estimated the extent of genetic assortative mating (GAM) by regressing *PGS*_*m*1_ on *PGS*_*m*2_:

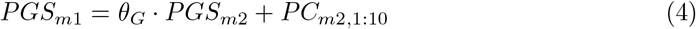

### Haplotype-based genetic assortative mating calculations

We estimated haplotype-based GAM by regressing the PGS calculated on one parental haplotype against the PGS of the other parental haplotype:

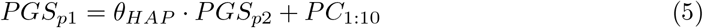

where *θ*_*HAP*_ represents the GAM estimate from offsprings only, and *PC*_1:10_ are the first 10 ancestry principal components included to account for population structure.

For some individuals, one or more chromosomes were missing their parental haplotype sets. This occurs when no IBD sharing is detected between an individual and their close relatives on a given chromosome, making inter-chromosomal phasing impossible for that chromosome^25^. To impute the resulting missing values in the PGS we used the mean PGS for the corresponding chromosome, computed across all individuals in the rest of the cohort. We verified that GAM estimates obtained with this imputation were highly consistent with estimates derived from the subset of individuals with complete parental haplotype data (*r*^2^ = 0.74; *y* = 1.01*x* + 0, RMSE=0.0043, Supplementary Figure 14).

### Gametic Phase Disequilibrium (GPD)-based genetic assortative mating estimation

We computed GPD-based GAM estimates following the method proposed by Yengo *et al*.^7^:

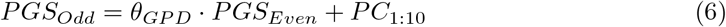

where *PGS*_*Odd*_ and *PGS*_*Even*_ are polygenic scores computed using SNPs located on odd- and even-numbered chromosomes, respectively. The GAM estimate 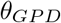 corresponds to the regression coefficient of *PGS*_*Even*_ on *PGS*_*Odd*_, adjusted for the first 10 principal components to account for population stratification.

To ensure a fair comparison between the GPD-based and our haplotype-based approaches, we restricted the GPD analysis to the subset of individuals for whom inter-chromosomal phasing was available, so that both methods were applied to the same sample set.

We note, however, that the original GPD-based implementation differs from ours in two important respects: (i) it was applied to a sample of unrelated individuals, whereas our sample included some related individuals; and (ii) it used PCs computed separately from odd or even chromosomes, while our version used genome-wide PCs. We therefore validated that both implementation produced consistent results (*r*^2^ = 0.75, *y* = 1.12*x*+0, *RMSE* = 0.0033, Supplementary Figure 15A, Supplementary Note 2).

## Data availability

### Code availability

The inter-chromosomal phasing pipeline is available at https://github.com/RJHFMSTR/THORIN^39^.

## Acknowledgments

We thank the participants of all biobanks for sharing their data. The Estonian Biobank data analysis was carried out in part in the High-Performance Computing Center of University of Tartu, Estonia. The UK Biobank data analysis was carried out in part in the High-Performance Computing Center of University of Lausanne, Switzerland. This research has been conducted using the UK Biobank Application Number 16389 and funded by the Swiss National Science Foundation (SNSF) project grant SNSF 310030-189147, 315230-219587. We thank Alexandre Reymond and Anthony Herzig for their valuable insights and discussions and Lili Milani for her assistance with the Estonian Biobank data access and processing.

## Ethics statement

The activities of the EstBB are regulated by the Human Genes Research Act, which was adopted in 2000 specifically for the operations of the EstBB. Individual level data analysis in the EstBB was carried out under ethical approval 1.1-12/295 from the Estonian Committee on Bioethics and Human Research (Estonian Ministry of Social Affairs), using data according to release application nr T38 from the Estonian Biobank.

## Authors information

### Authors contributions

R.J.H. and Z.K. designed the study, methods and statistical tests. R.J.H. and Z.K. wrote the manuscript. R.J.H. performed all data analyses. T.C assisted with the derivation and evaluation of the accuracy of polygenic scores. D.M., E.P. and G.M. assisted with the implementation of the GPD-based approach. The Estonian Biobank research team performed data collection, genotyping and QC of the Estonian Biobank genetic data. L.P. and R.M. contributed to data curation. L.P., D.M., R.M., N.B., L.K., T.S. contributed to the interpretation of the result. The project was supervised by Z.K. All authors reviewed and approved the final manuscript.

## Additional authors affiliations

### Estonian Biobank research team

Andres Metspalu^4^, Lili Milani^4^, Tõnu Esko^4^, Reedik Mägi^4^, Mari Nelis^4^ and Georgi Hudjashov^4^

## Supplementary Notes

### Supplementary Note 1 Estimating assortative mating strength from individual genomes

#### 1.1 Theoretical derivations

##### 1.1.1 Problem setting

Let 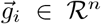 denote the centred (i.e. zero-mean) polygenic score (PRS) of a set of *n* individuals for a fixed trait on chromosome *i* computed based on the first parent’s haplotype (either mother of father) and 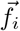 the same quantity when using the other haplotype. Let us assume that the parents of these individuals assorted for this trait and the correlation between the partners traits is *R*, i.e. the strength of assortment. Let the polygenic score based on the untransmitted alleles from the first parents on chromosome *i* be 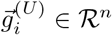 and the same quantity for the other parent is 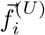. Thus, the PRS for the first parent based on both chromosome *i*s is 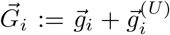 and the same (diploid)PRS for the other parent is 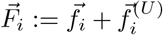. Let us define 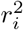 as the explained variance of the diploid polygenic score on chromosome *i* of the trait.

##### 1.1.2 Estimating assortative mating strength from intra-chromosome phased individuals

Our first aim is to estimate *R* only using the correlation between haploid PRSs in the offspring. As a first step we relate the correlation between parental PRSs and assortative mating strength for this trait.

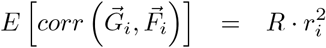

Thus, we can obtain an estimator for the assortative mating strength

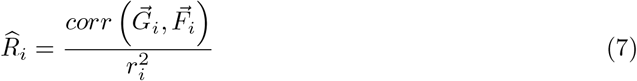

Now, we connect the haploid with diploid PRS correlations

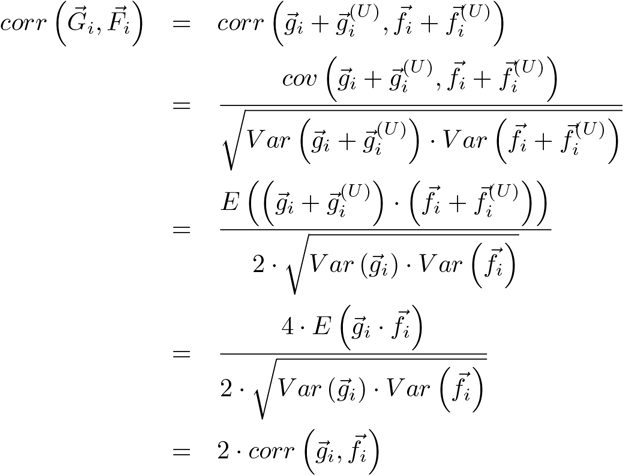

Therefore,

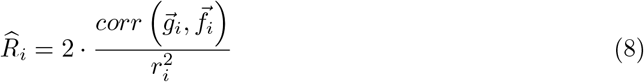

with variance

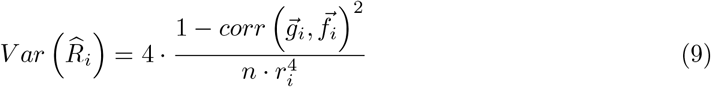

We can perform inverse-variance weighted meta-analysis of these estimates as follows

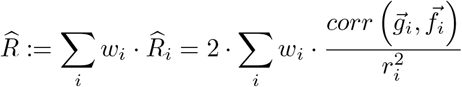

where

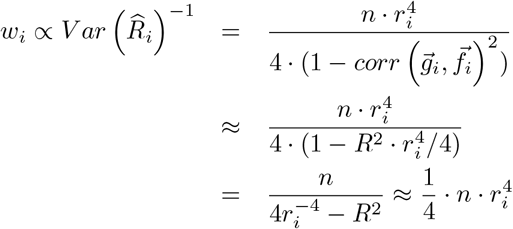

where the weights sum up to one, i.e.

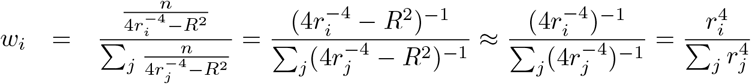

Therefore, the estimator simplifies to

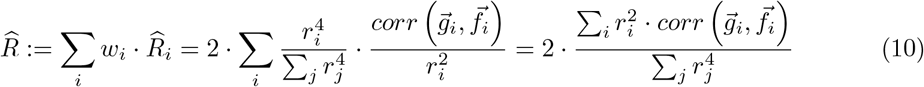

This allows us the computation of the variance of the estimator

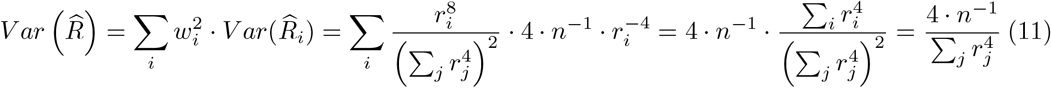

##### 1.1.3 Estimating assortative mating strength from inter-chromosome phased individuals

If we can do inter-chromosome phasing, the haploid PRS can be built for the whole genome instead of just one chromosome at a time. The whole genome PRS can be written as

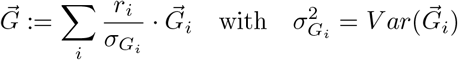

and

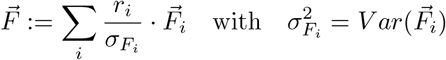

These PRSs explains 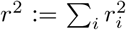 variance of the trait. In that situation, we have a better estimator for *R* from these haploid PRSs:

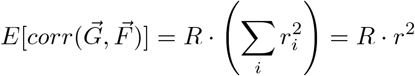

Let us now switch from diploid to haploid PRSs. As we have seen above, the diploid and haploid correlations can be related to each other as follows

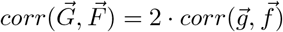

The haploid PRSs can be built as follows

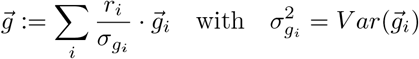

and

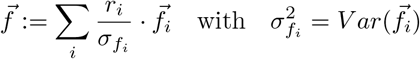

Thus the haploid PRS based estimator for assortment, *R* can be written as

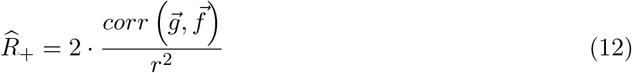

The variance of this estimator is thus

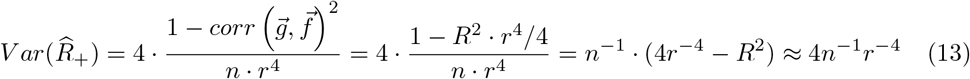

the last approximation is based on the fact that *r*^−4^ >> *r*^2^. We can compare this variance to the one obtained from the intra-chromosome phasing based one by assuming that the explained variance for chromosome *i* is proportional to the length of chromosome *i*, i.e.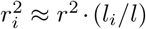, where *l*_*i*_ is the length of chromosome *i* and 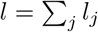 Therefore, the variance of the intra-chromosome phased based estimator variance is.

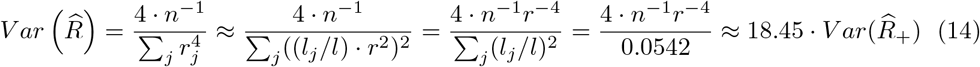

##### 1.1.4 Summary

This means the number of individuals with inter-chromosome phased PGS yields the same accuracy as 18 times intra-chromsomally phased individuals. For example in the UK Biobank, we can intra-chromosomally phase all 500,000 individuals, but the 240,000 for whom intra-chromosomal phasing is available are worth *>* 4 million intra-chromosomally phased individuals. If we take human height as example the value of *R* ≈ 0.3 and *r*^2^ ≈ 0.25 and *n* = 240, 000, thus the test statistic is 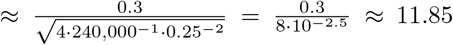 Thus, assortment strength is well detectable.

#### 1.2 Real data validation

We validated our haplotype-based GAM estimation approach using real data from the Estonian Biobank, which contains a large number of parent-offspring trios (N=14,063), corresponding to 10,618 unique mate pairs. This dataset enables a direct comparison of GAM estimates obtained from different methods.

We first computed GAM directly from the genotypes of mate pairs, which we consider the ground truth. We then applied our haplotype-based GAM method to the corresponding 10,618 offspring, retaining only one offspring per parent pair to avoid biasing estimates by including siblings (who represent the same parental mating event).

Our results show that the haplotype-based GAM estimates derived from offspring were highly concordant with those obtained directly from their parents (*r*^2^ = 0.559, *β* = 0.49, Supplementary Figure 2), demonstrating that our approach accurately recovers recent assortative mating patterns in the parental generation. This concordance supports the validity of our method as a scalable alternative to mate-pair-based analyses in large cohorts lacking explicit couple information.

## Supplementary Note 2 Variation in GPD-based GAM calculation

While we implemented the GPD-based approach on the same sample set as our haplotype-based approach, we note, that the original GPD-based implementation^7^ differs from ours in two important respects: (i) it was applied to a sample of unrelated individuals, whereas our sample included some related individuals; and (ii) it used PCs computed separately from odd or even chromosomes, while our version used genome-wide PCs.

To evaluate the impact of these differences, we implemented and compared two versions of the GPD-based approach in the Estonian Biobank: variation 1, our implementation, and variation 2, which followed the original protocol as closely as possible. We performed this comparison in the Estonian Biobank specifically, as it allowed us to ensure that no individuals used to derive the PGS overlapped with those in the GPD analysis.

To identify the largest set of unrelated individuals in the Estonian Biobank, we first used the KING software v2.2.4^33^ to compute genetic relatedness. We then iteratively removed the individual with the largest number of close relatives (up to the 4^*th*^ degree), until no relatedness remained, for a final set of 48,155 individuals.

We compared variation 1 to variation 2, and found highly concordant results (*r*^2^ = 0.75, *y* = 1.12*x* + 0, *RMSE* = 0.0033, Supplementary Figure 15A). Notably, variation 1 identified significant GAM for two additional traits (Supplementary Figure 15A), likely due to the increased sample size afforded by including related individuals. When restricted to the same sample set (i.e only the use of PCs varies), it was even more concordant (*r*^2^ = 0.95, *y* = 0.94*x* + 0, *RMSE* = 0.0014).

Finally, we compared variation 1 and variation 2 against mate-pairs estimates, and both implementations showed high concordance (*r*^2^ = 0.48 and 0.41); *y* = 0.19*x*+0 and *y* = 0.23*x*+0; RMSE = 0.015 and 0.014, for variation 1 and variation 2, respectively; Supplementary Figure 15B,C).

## Supplementary Figures

**Supplementary Fig. 1.**
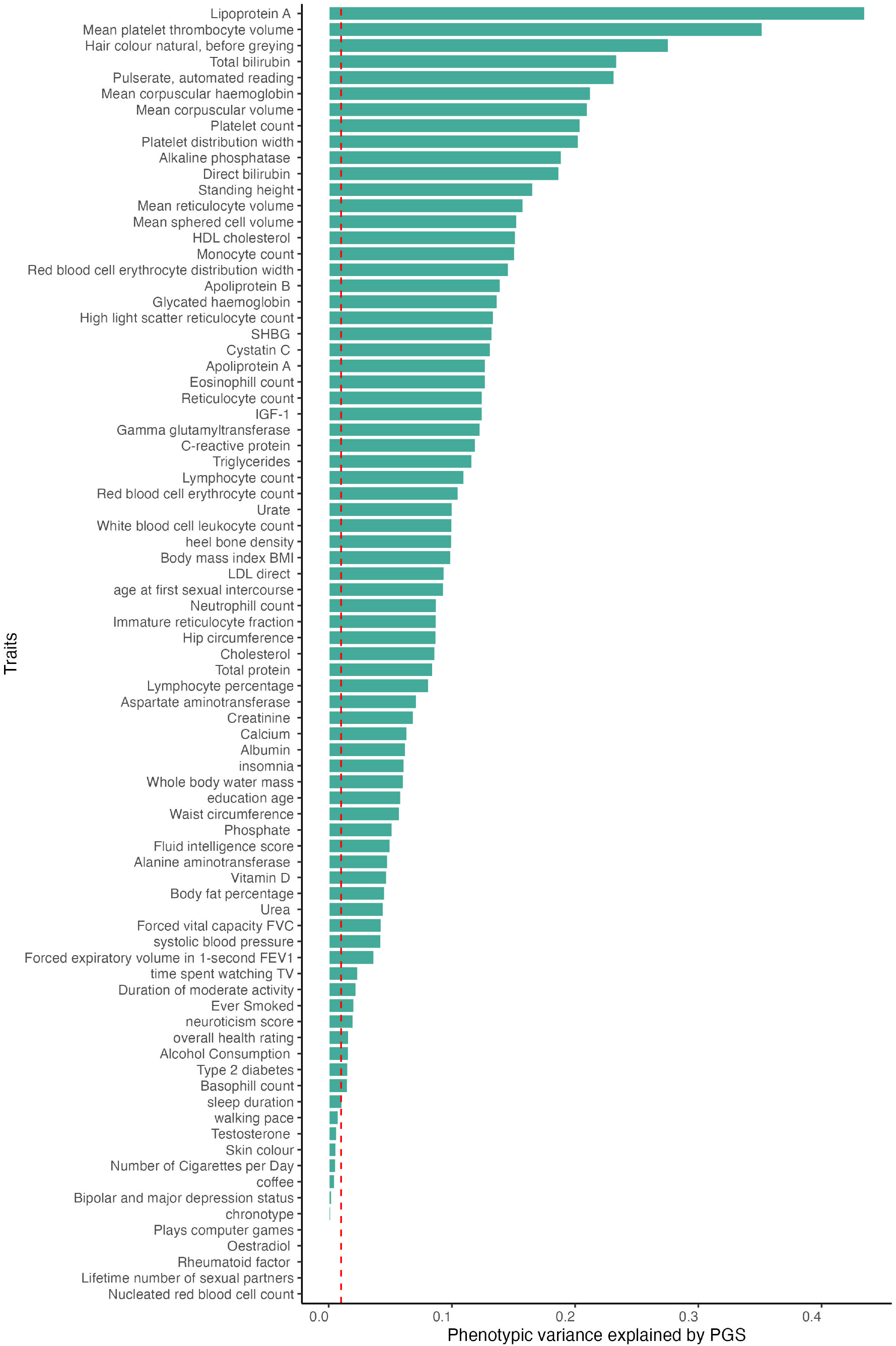
Phenotypic variance explained by PGS in the UK Biobank. Phenotypic variance explained by the PGS (*r*^2^, x-axis) across the 81 selected traits (y-axis). *r*^2^ was computed in an out-of-sample set of 245,884 white British UK Biobank individuals (i.e those not used for association testing). Red dashed line shows *r*^2^ = 0.01. We kept only the 69 traits with *r*^2^ *>* 0.01 for downstream analysis.

**Supplementary Fig. 2.**
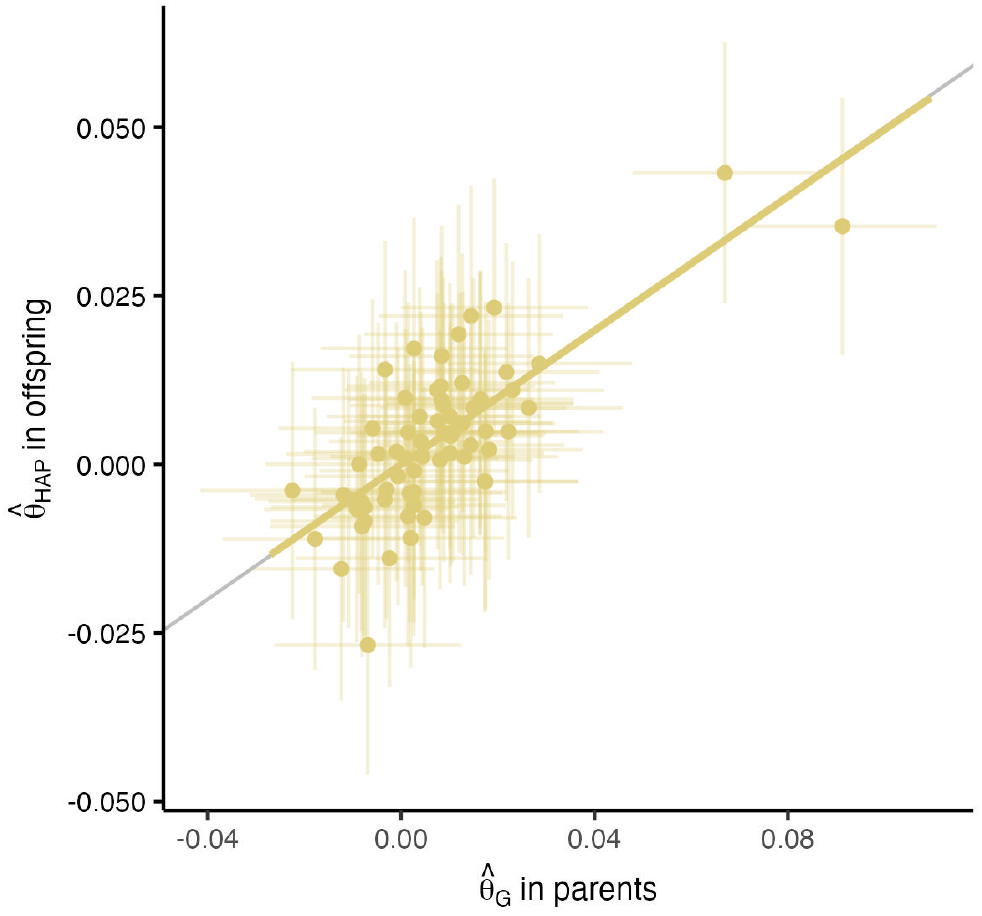
Method validation in the Estonian Biobank. GAM computed with our haplotype-based approach in offspring (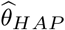, y-axis) *vs*. computed from parents 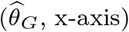 in the Estonian Biobank. Each dot show 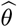 values for a different trait, with error bars indicating 95% confidence intervals computed as 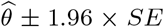. Grey line shows the expected line *y* = 0.5*x* + 0. Yellow line shows the regression line *y* = 0.49*x* + 0.

**Supplementary Fig. 3.**
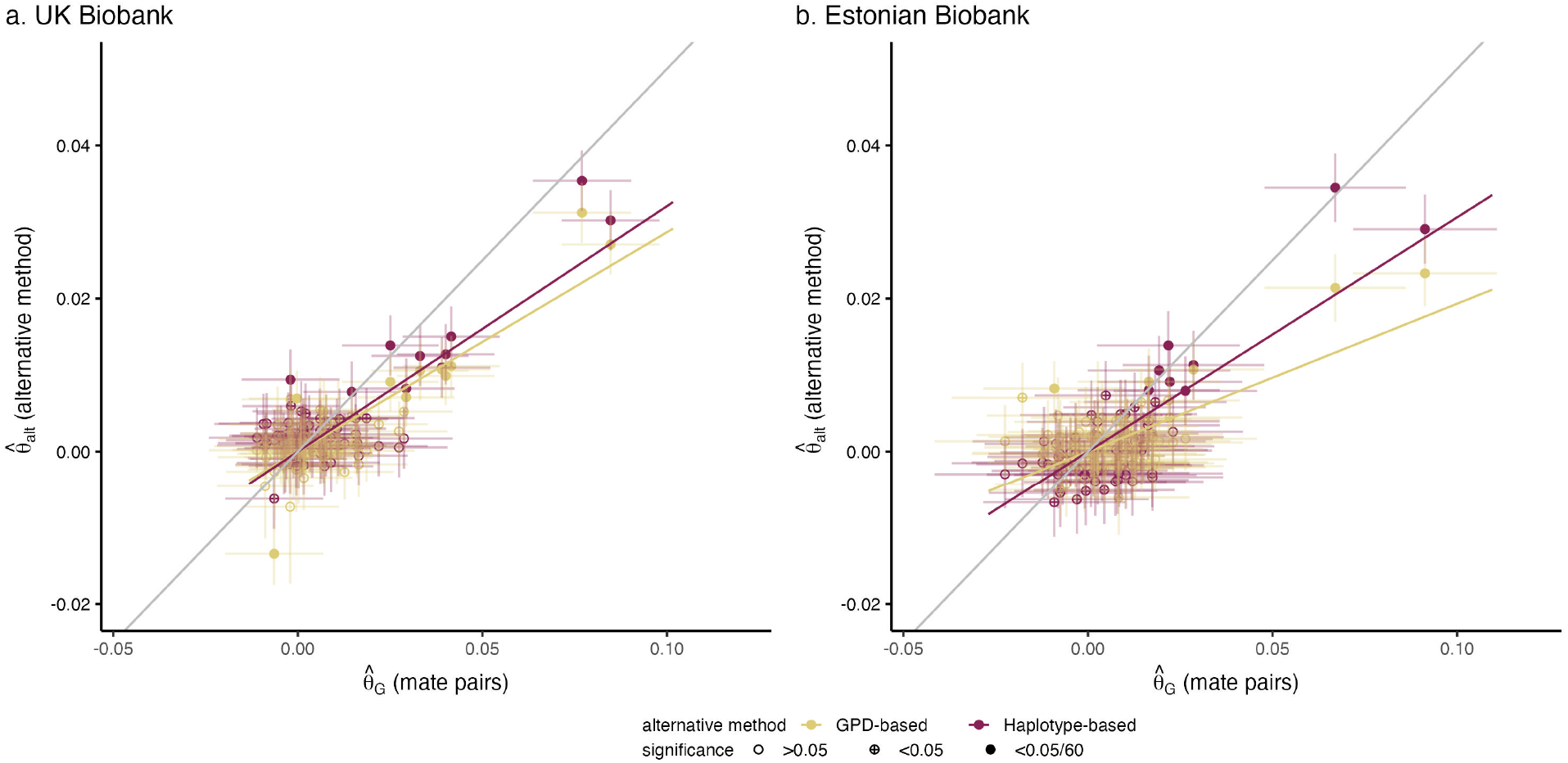
GAM estimates performance. GAM computed from mate-pairs 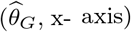 *vs*. GAM computed using the haplotype-based or GPD-based approach (y-axis; purple indicates haplotype-based and yellow indicates GPD-based) in the UK (**a**) and Estonian (**b**) Biobank. Each dot show 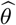 values for a different trait, with error bars indicating 95% confidence intervals computed as 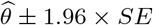. Grey line shows the expected line *y* = 0.5*x* + 0. Purple and yellow lines show the regression lines for the haplotype-based and GPD-based approaches, respectively.

**Supplementary Fig. 4.**
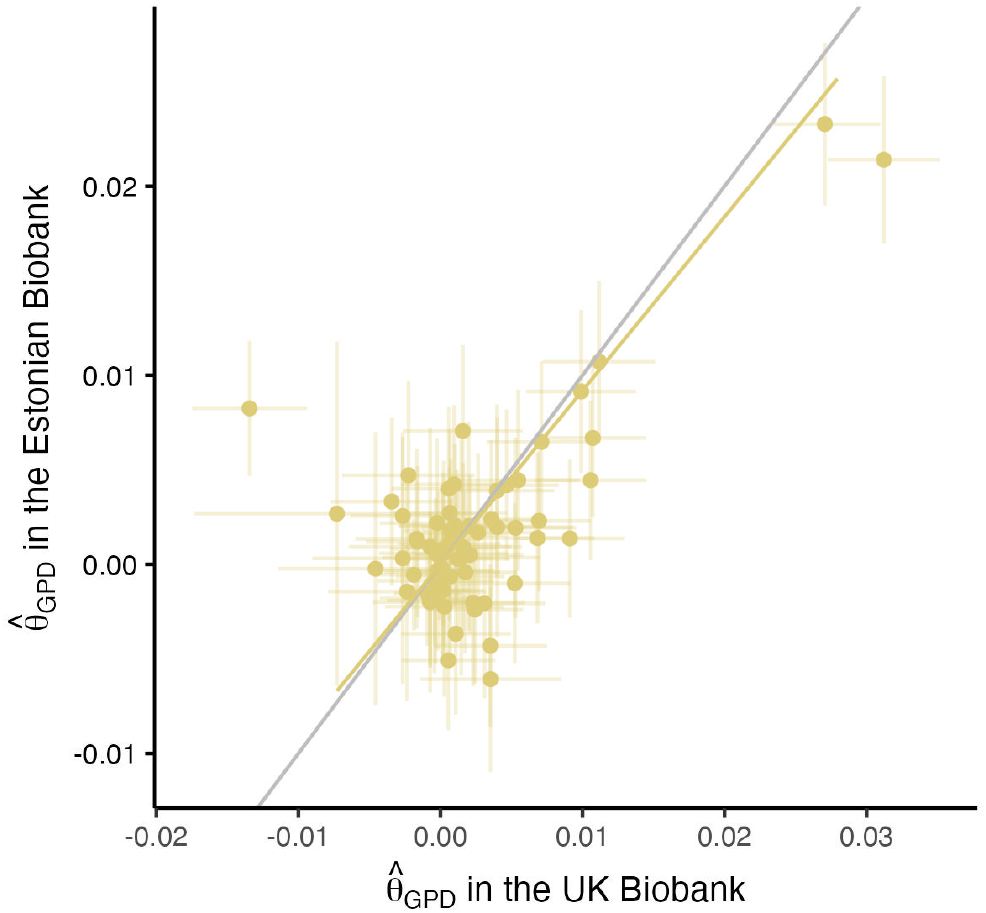
GAM estimates comparison between the UK and the Estonian cohorts. GAM computed with the GPD-based approach 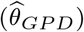 in the UK (x-axis) *vs*. Estonian Biobank (y-axis). Each dot show the 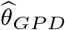 for a different trait, with error bars indicating 95% confidence intervals computed as 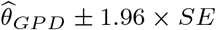. Grey line shows the identity line. Yellow line shows the regression line.

**Supplementary Fig. 5.**
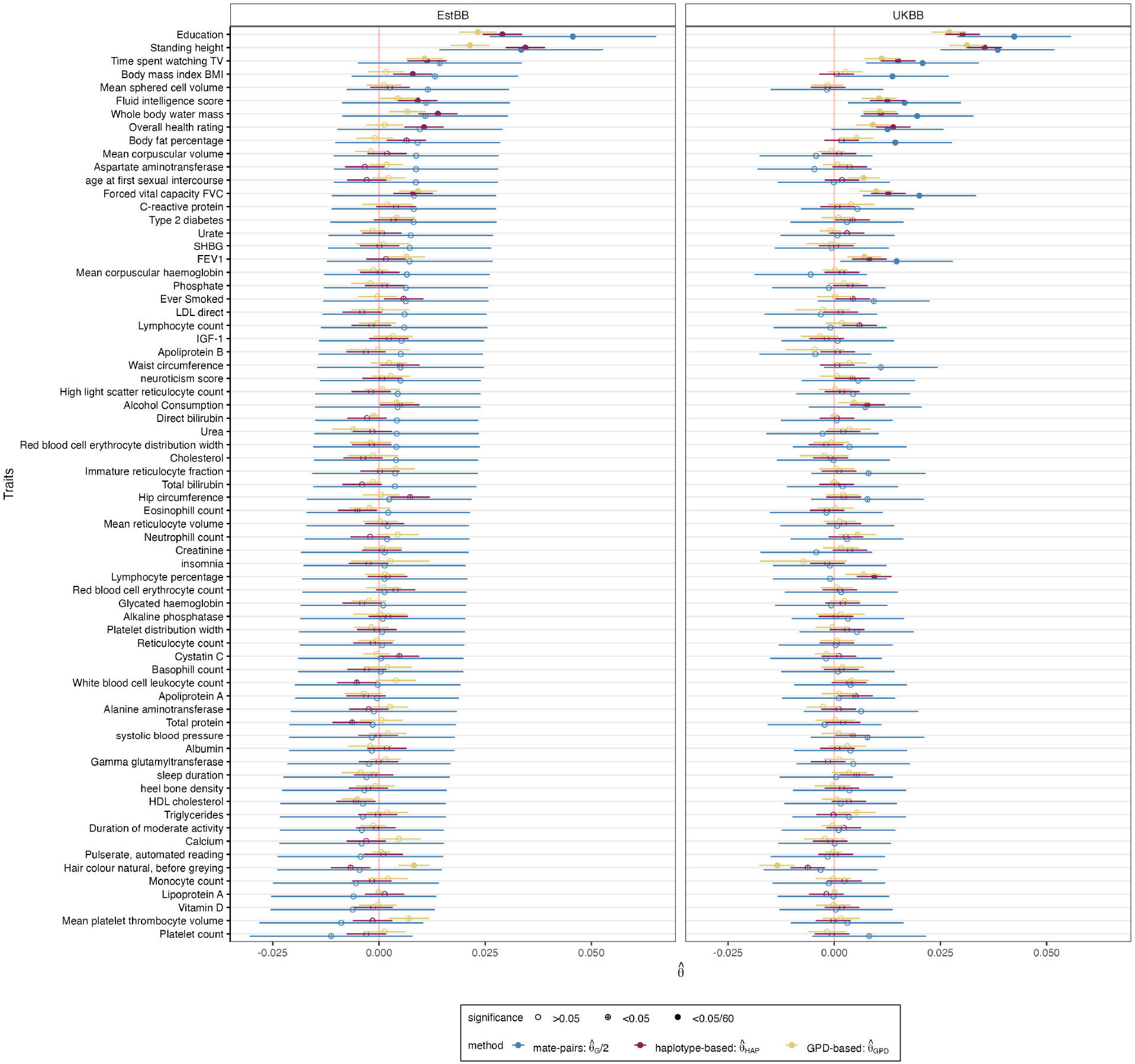
GAM estimates in the UK and the Estonian cohorts. GAM estimates (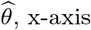 blue for mate-pair 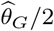 (rescaled for comparability); purple for haplotype-based 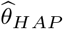; yellow for GPD-based 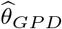) across the 69 selected traits (y-axis) with PGS *r*^2^ *>* 0.01. Shapes indicate non-significant (empty circle), nominally significant (crossed circle) and Bonferroni significant (full circle) traits. Error bars show 95% confidence intervals computed as 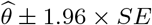.

**Supplementary Fig. 6.**
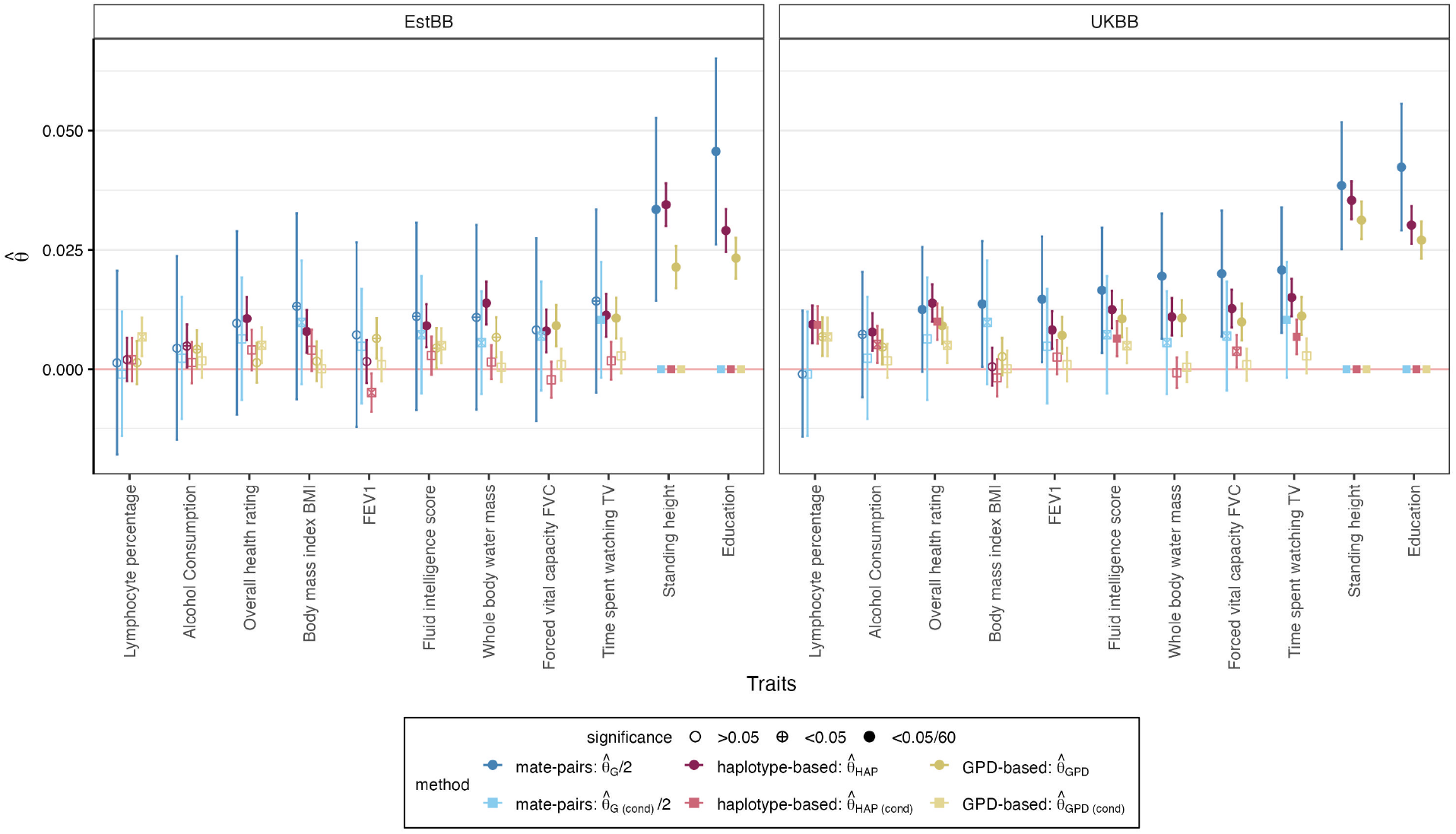
Conditional GAM estimates in the UK and the Estonian cohorts. GAM estimates (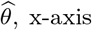; blue for mate-pair 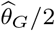 (rescaled for comparability); purple for haplotype-based 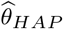; yellow for GPD-based 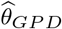) across traits (y-axis) with PGS *r*^2^ *>* 0.01. Dark and light colors indicate original estimates and estimates when conditioned on standing height and educational attainment, respectively. Shapes indicate non-significant (empty circle), nominally significant (crossed circle) and Bonferroni significant (full circle) traits. Error bars show 95% confidence intervals computed as 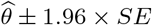.

**Supplementary Fig. 7.**
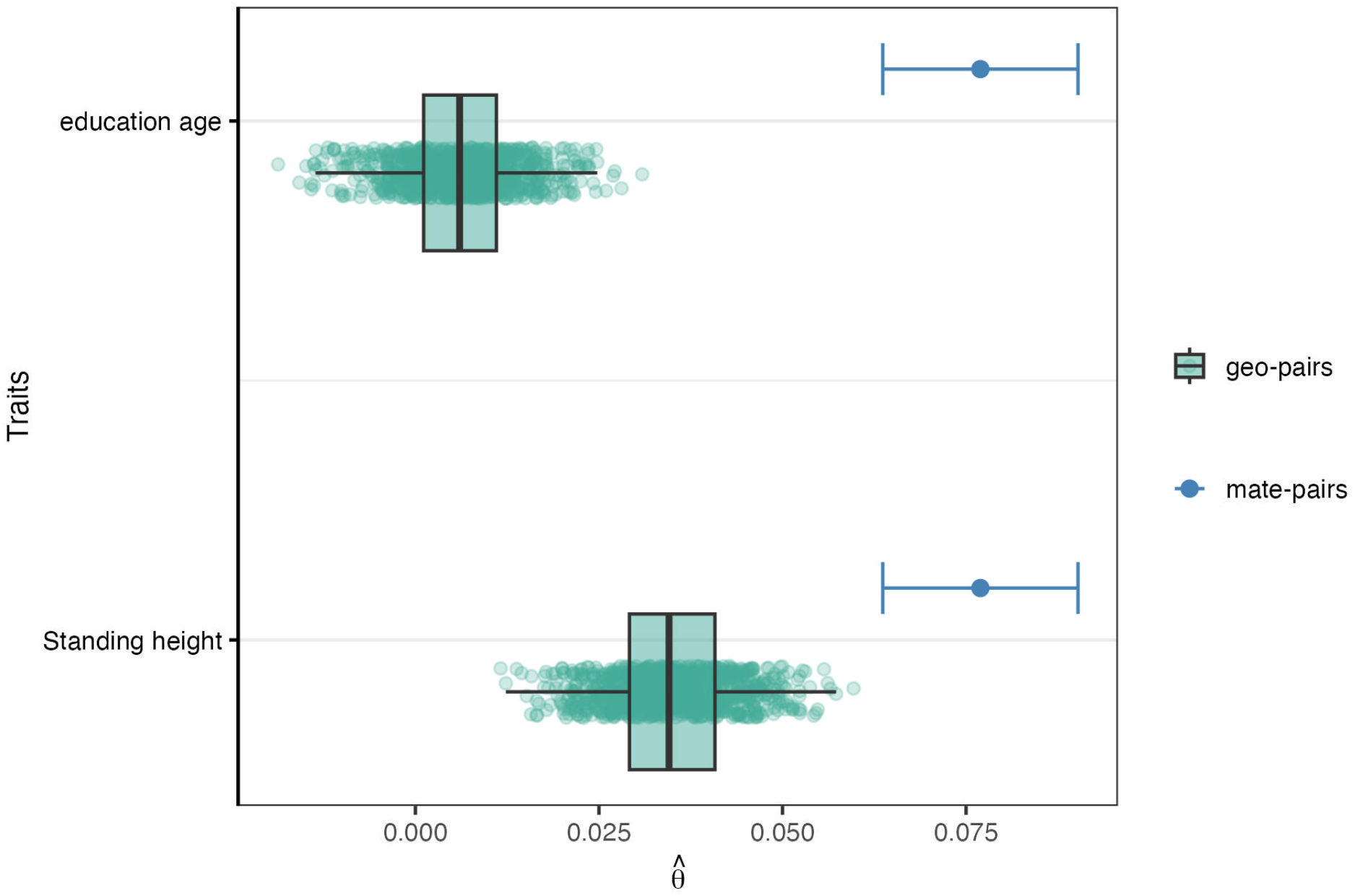
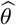 for geo-pairs based on spatial proximity. GAM (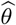, x-axis) estimated from mate-pairs (blue; 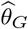) and from geo-pairs (green; 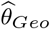) for education attainment and standing height (y-axis). For mate-pairs, error bars show 95% confidence intervals computed as 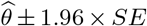. For geo-pairs, boxes indicate the interquartile range (IQR), with the bottom and top of the box representing the 25th (Q1) and 75th (Q3) percentiles, respectively. The horizontal line within the box represents the median (50th percentile). Whiskers extend to the smallest and largest values within *Q*1−1.5 *× IQR* and *Q*3 + 1.5 *× IQR*. Each dot represent the 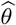 computed from 20,000 randomly sampled geo-pairs (sampling repeated 1,000 times).

**Supplementary Fig. 8.**
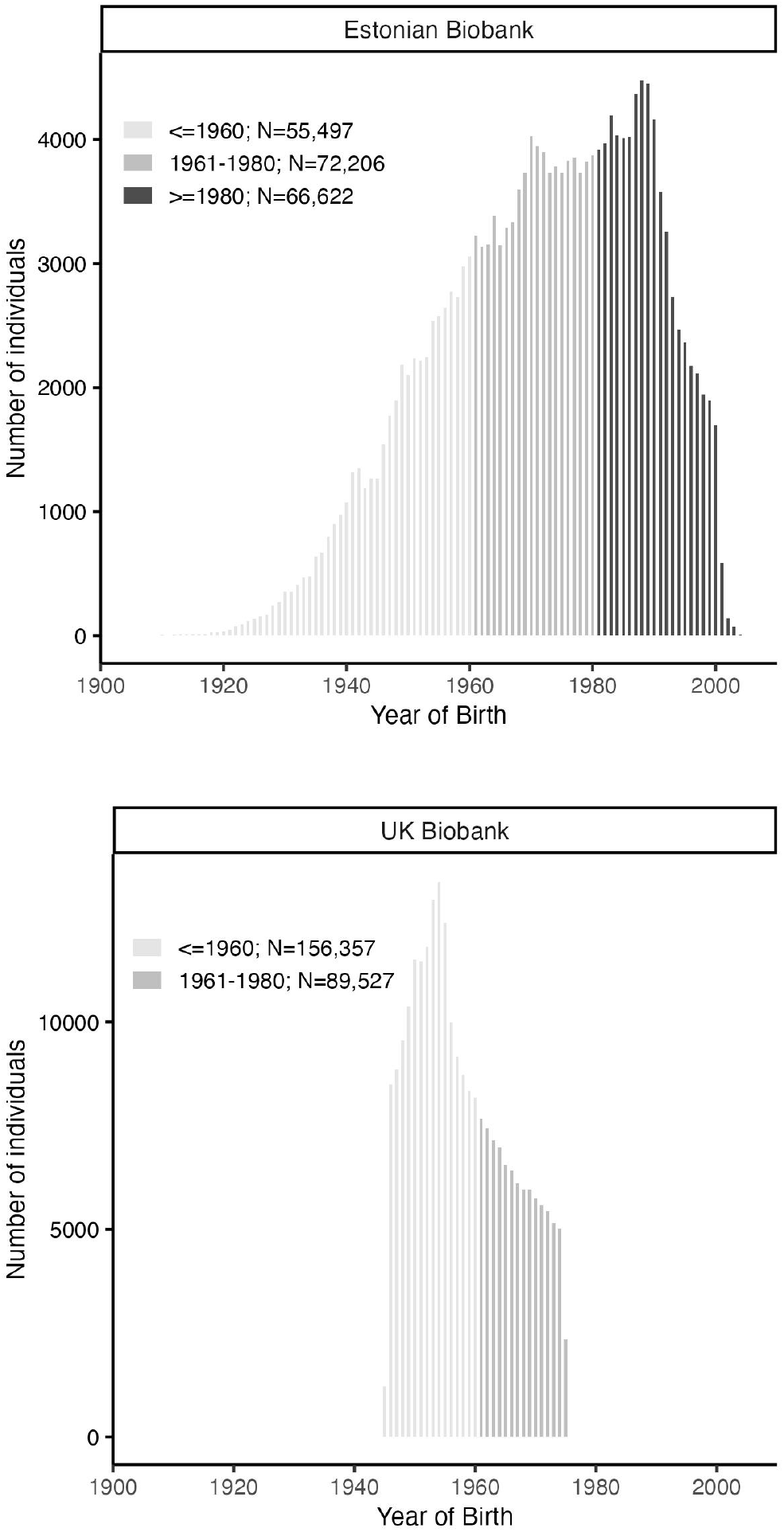
Year of Birth (YoB) bins and distribution in the UK and Estonian Biobanks. Distribution of the number of individuals (y-axis) per YoB (x-axis) in the Estonian (top) and UK (bottom) Biobanks. Each bar represent a YoB. Grey colors indicate the bins used in this study (*Y oB <*= 1960; 1961 *< Y ob <*= 1980; *Y oB >* 1980)

**Supplementary Fig. 9.**
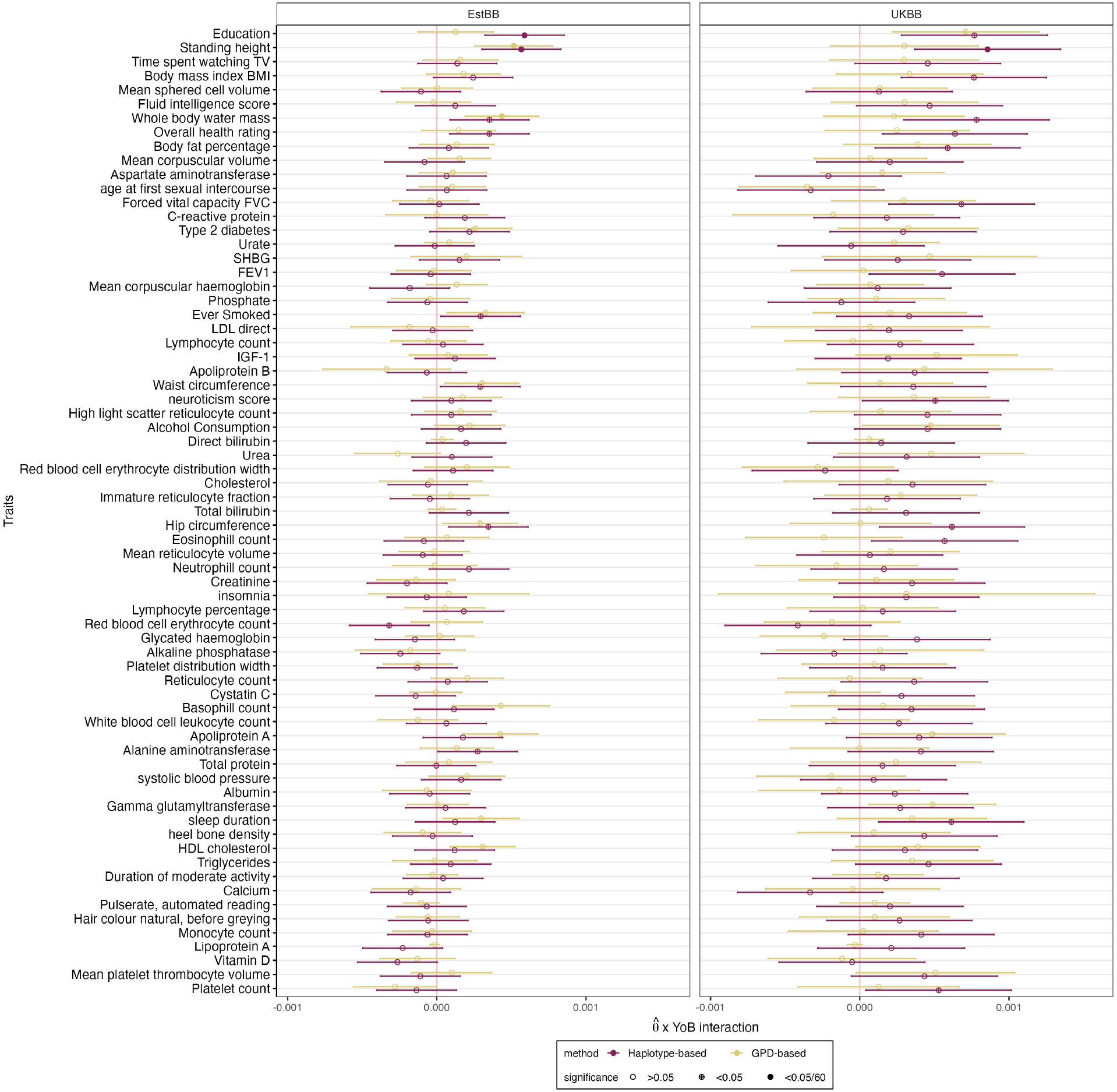
Year of Birth (YoB) interaction effect on 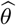. YoB interaction effect (x-axis) on 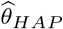 (purple) and 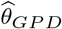 (yellow) across the 69 traits (y-axis). Shapes indicate non-significant (empty circle), nominally significant (crossed circle) and Bonferroni significant (full circle) traits. Error bars show 95% confidence intervals computed as 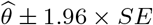.

**Supplementary Fig. 10.**
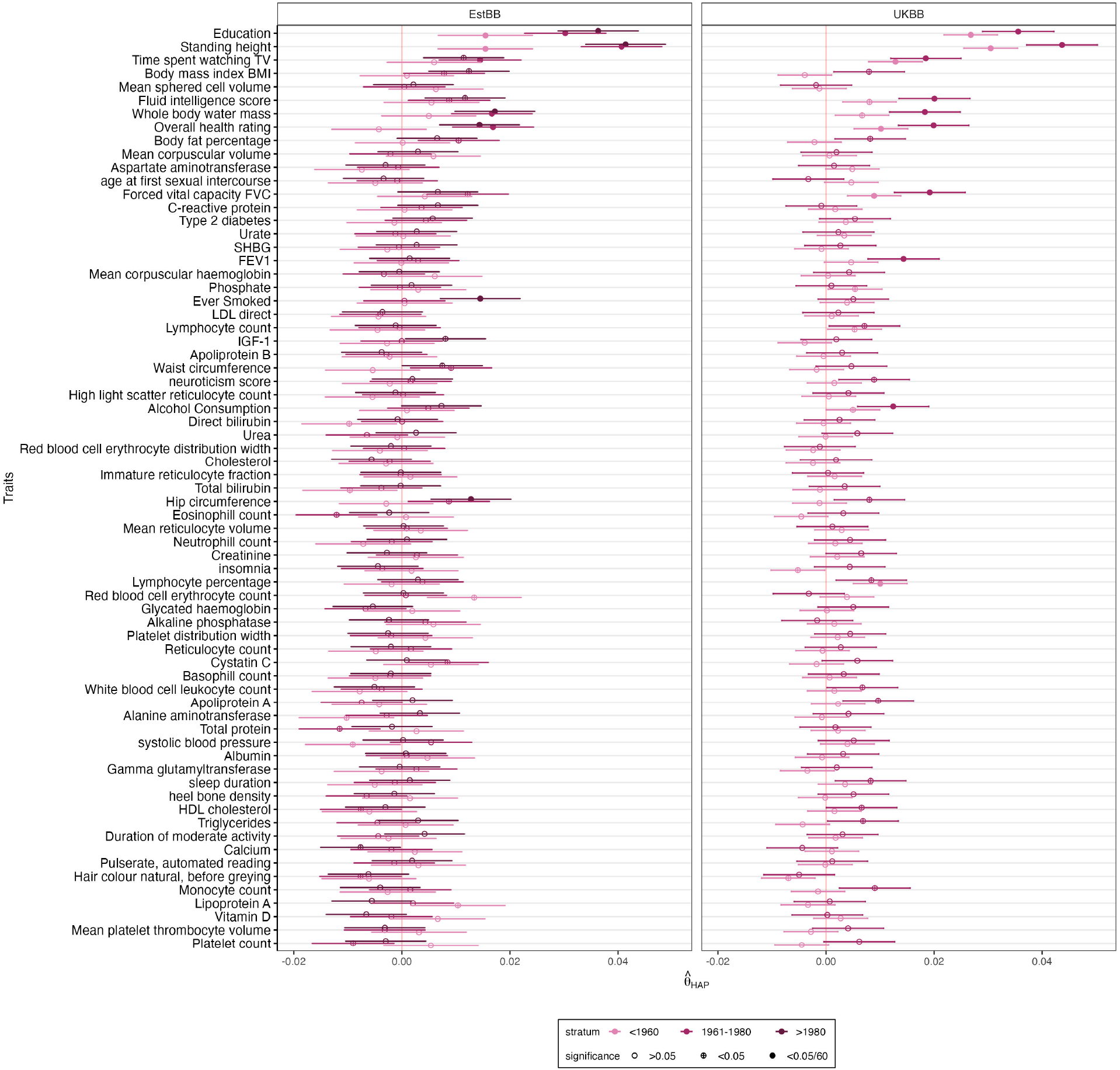
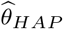 per Year of Birth (YoB) bins. Haplotype-based GAM (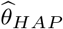, x-axis) per YoB bins across the 69 traits (y-axis). Colors represent YoB bins, with darker colors indicating more recent generations. Shapes indicate non-significant (empty circle), nominally significant (crossed circle) and Bonferroni significant (full circle) traits. Error bars show 95% confidence intervals computed as 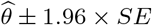.

**Supplementary Fig. 11.**
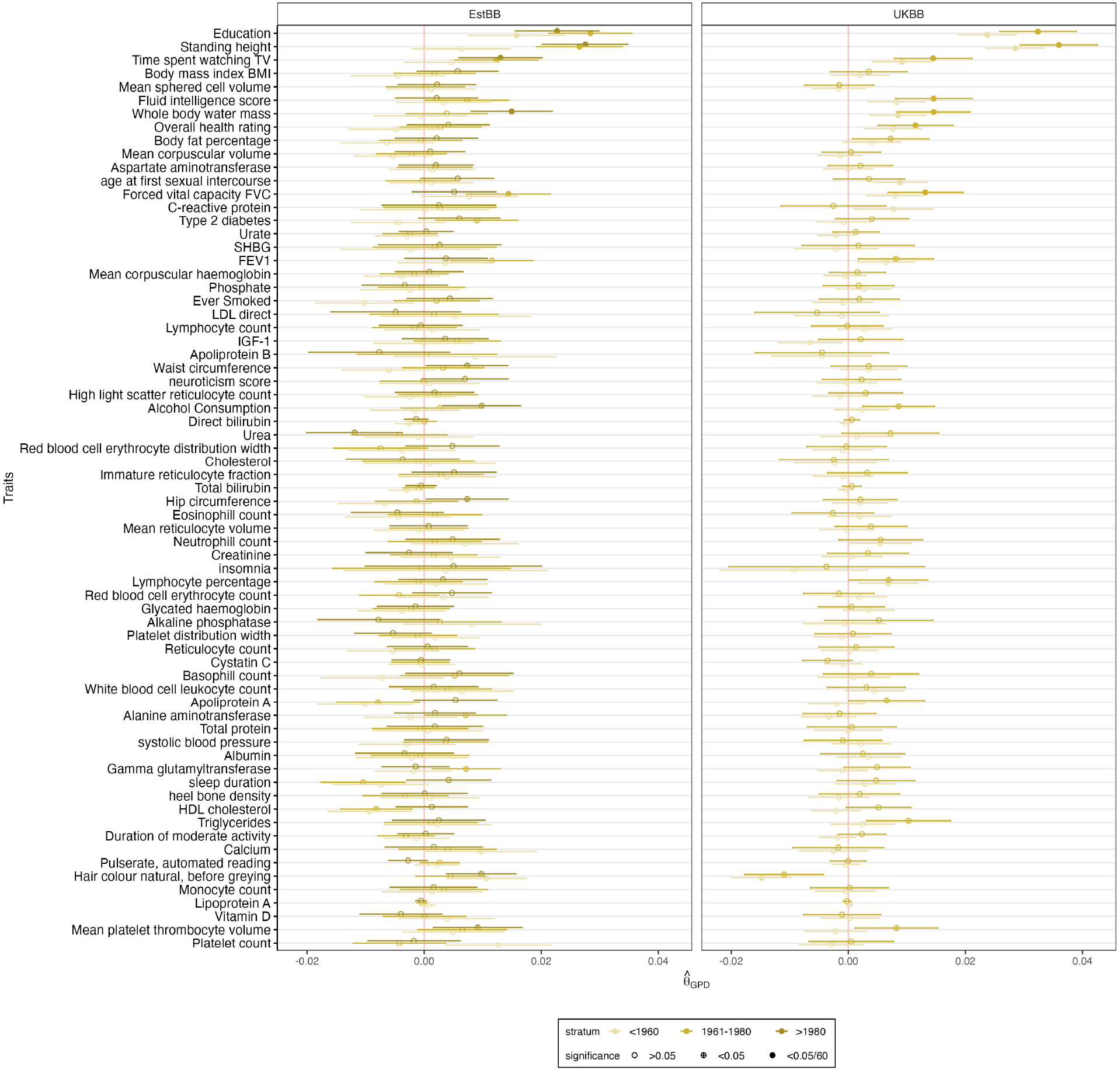
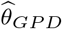 per Year of Birth (YoB) bins. GPD-based GAM (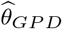, x-axis) per YoB bins across the 69 traits (y-axis). Dot colors represent YoB bins, with darker colors indicating more recent generations. Shapes indicate non-significant (empty circle), nominally significant (crossed circle) and Bonferroni significant (full circle) traits. Error bars show 95% confidence intervals computed as 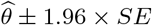.

**Supplementary Fig. 12.**
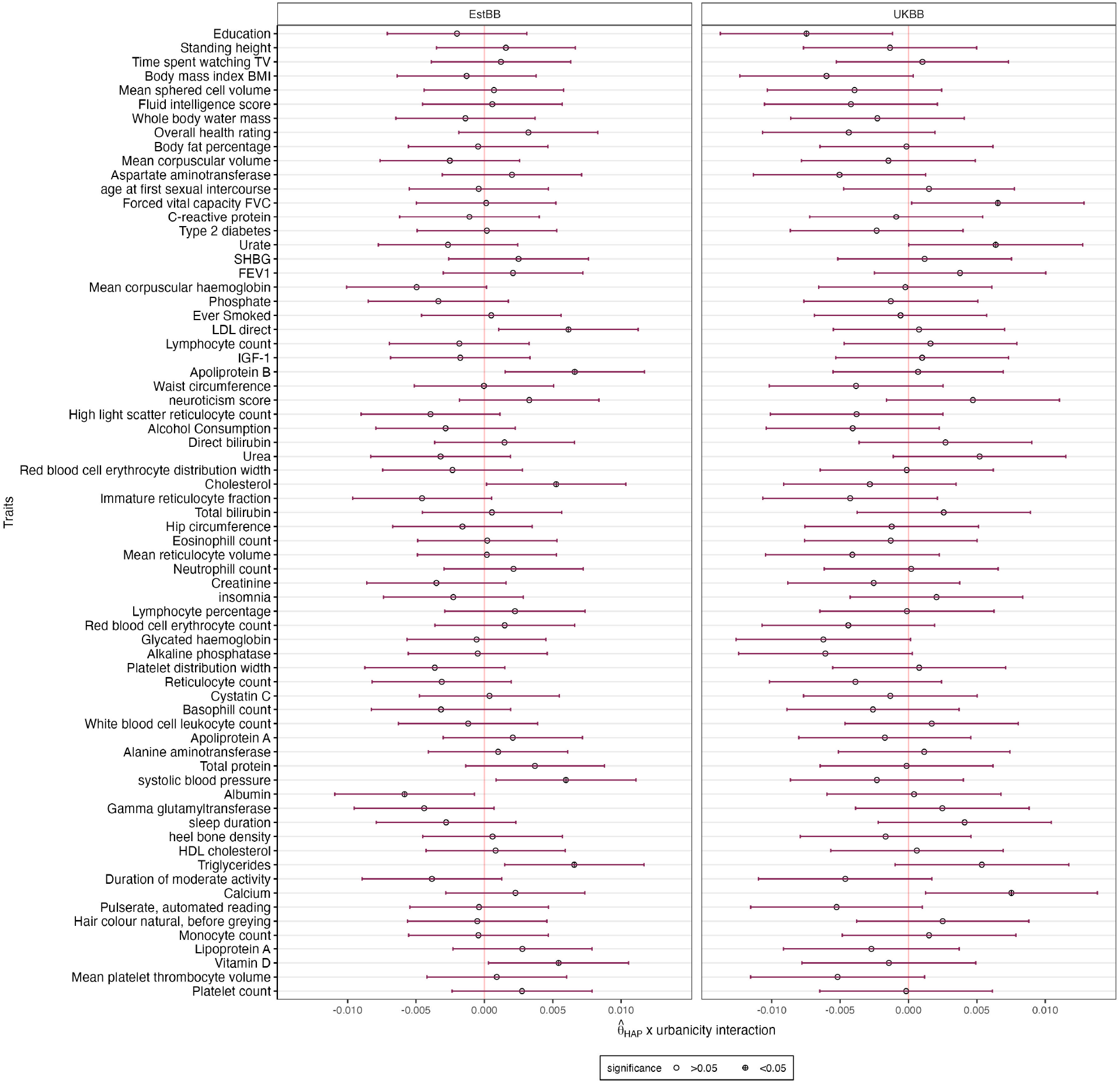
Urbanicity interaction effect on 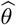. Urbanicity interaction effect (x-axis) on 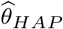 (purple) and 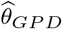 (yellow) across the 69 traits (y-axis). Shapes indicate non-significant (empty circle), nominally significant (crossed circle) and Bonferroni significant (full circle) traits. Error bars show 95% confidence intervals computed as 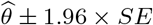.

**Supplementary Fig. 13.**
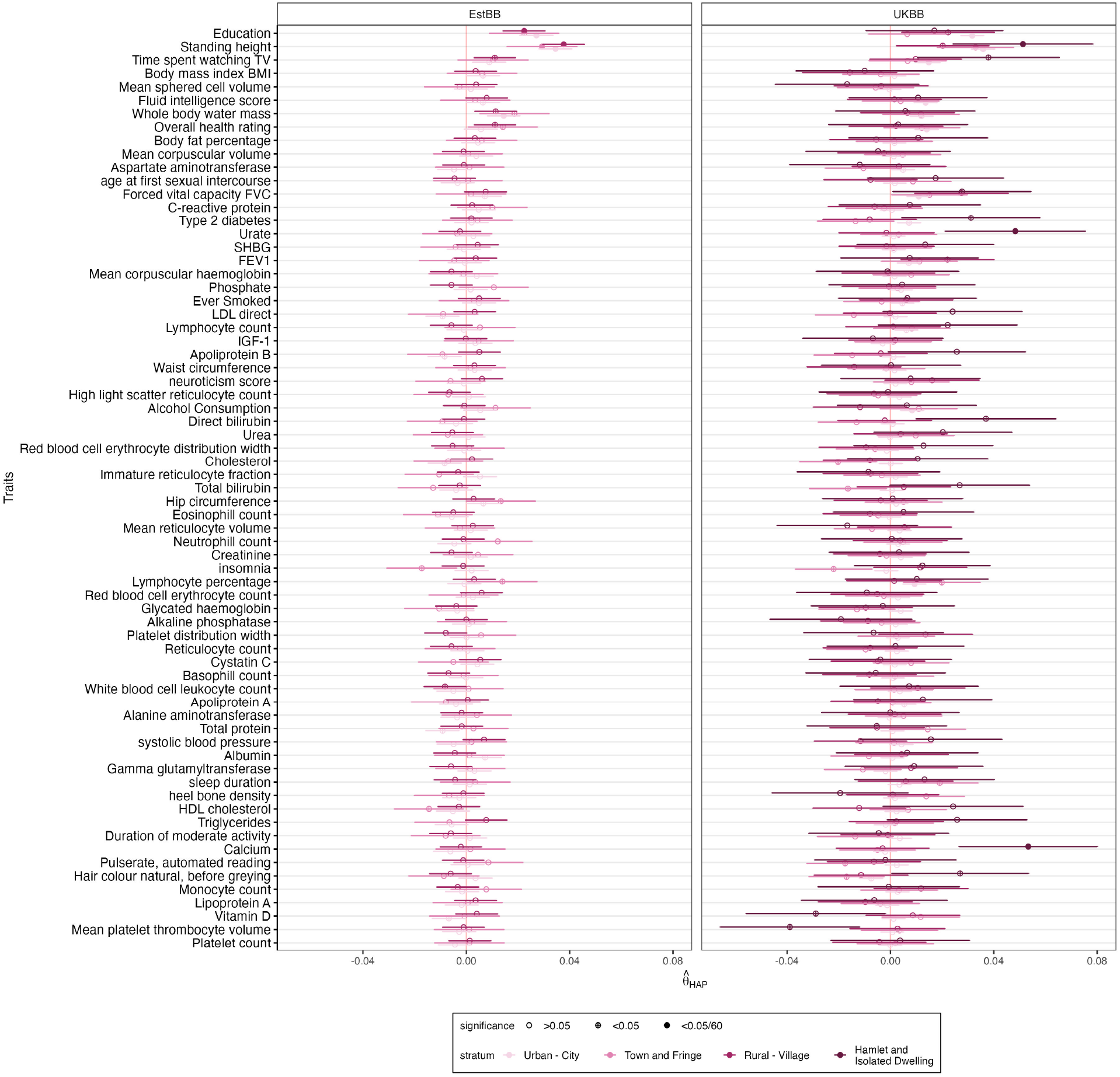
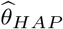 per settlement type. Haplotype-based GAM (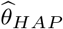, x-axis) per settlement type across the 69 traits (y-axis). Dot colors represent settlement types, with darker colors indicating more rural areas. Shapes indicate non-significant (empty circle), nominally significant (crossed circle) and Bonferroni significant (full circle) traits. Error bars show 95% confidence intervals computed as 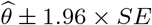.

**Supplementary Fig. 14.**
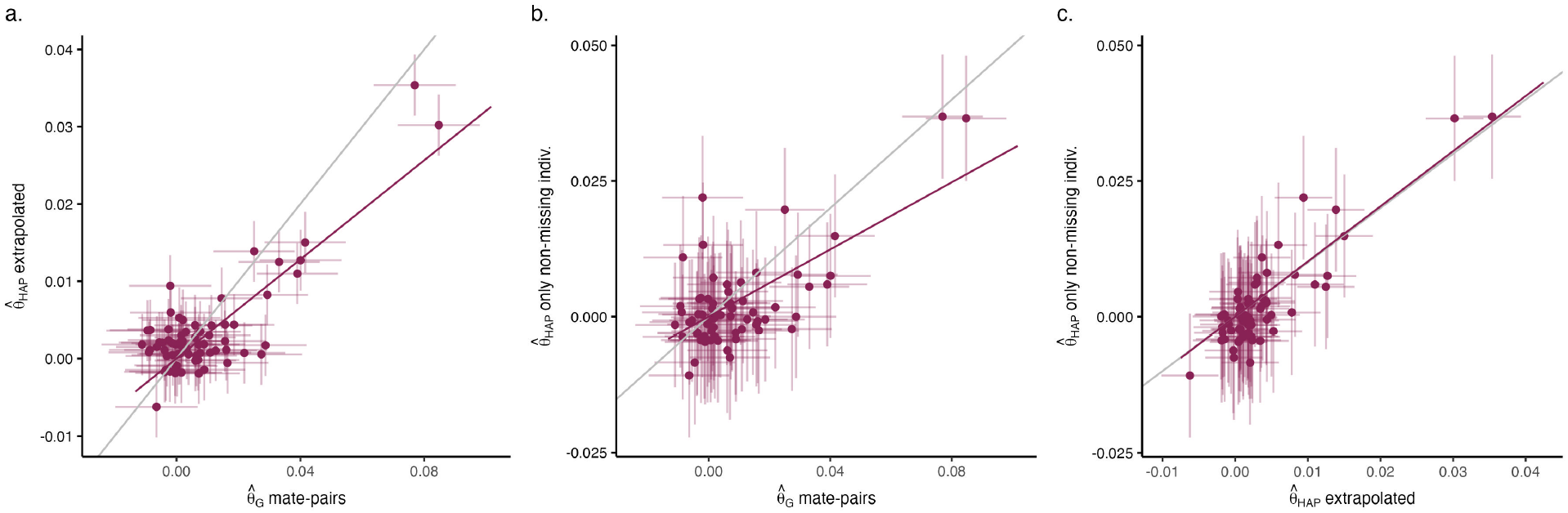
Haplotype-based GAM method comparison. **a)** GAM computed from mate-pairs (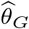, x-axis) *vs*. GAM computed using our haplotype-based approach (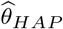, y-axis), where the PGS from missing chromosome is extrapolated from the population, as implemented broadly in our study. **b)** GAM computed from mate-pairs (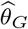, x-axis) *vs*. GAM computed using our haplotype-based approach (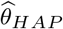, y-axis), where we used only individuals with non-missing PGS. **c)** 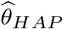, where the PGS from missing chromosome is extrapolated from the population, as implemented in our study (x-axis) *vs*. 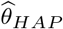, where we used only individuals with non-missing PGS (y-axis). For **a, b** and **c**, each dot show the 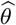 for a different trait, with error bars indicating 95% confidence intervals computed as 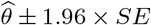. Grey lines shows the expected regression line (*y* = 0.5*x* + 0 for **a** and **b**; *y* = 1*x* + 0 for **c**). Purple line shows the observed regression lines.

**Supplementary Fig. 15.**
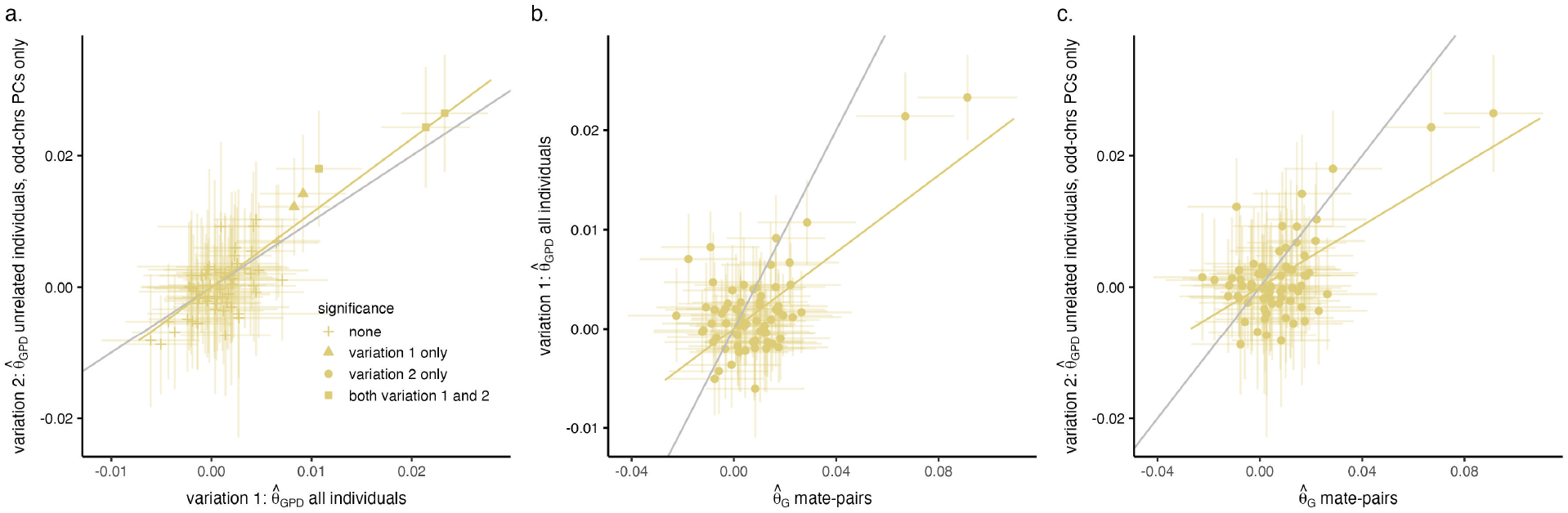
GPD-based GAM method comparison. **a)** 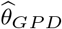 on all individuals with available interchromosomal phasing, as implemented in our study (variation 1, x-axis) *vs*. 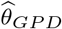 using the largest set of unrelated individuals and PCs derived from odd-numbered chromosomes only, as original implemented^7^ (variation 2, y-axis). **b)** 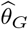 computed from mate-pairs (x-axis) *vs*. 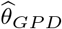 variation 1 (y-axis). **c)** 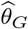 computed from mate-pairs (x-axis) *vs*. 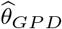 variation 2 (y-axis). For **a, b** and **c**, each dot show the 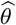 for a different trait, with error bars indicating 95% confidence intervals computed as 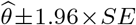. Grey lines shows the expected regression line (*y* = 1*x*+0 for **a**; *y* = 0.5*x*+0 for **b** and **c**). Yellow line shows the observed regression lines. In **a**, shapes indicate which GPD-based implementation(s) found the trait significant.

## Supplementary Tables

**Supplementary Table 1.**
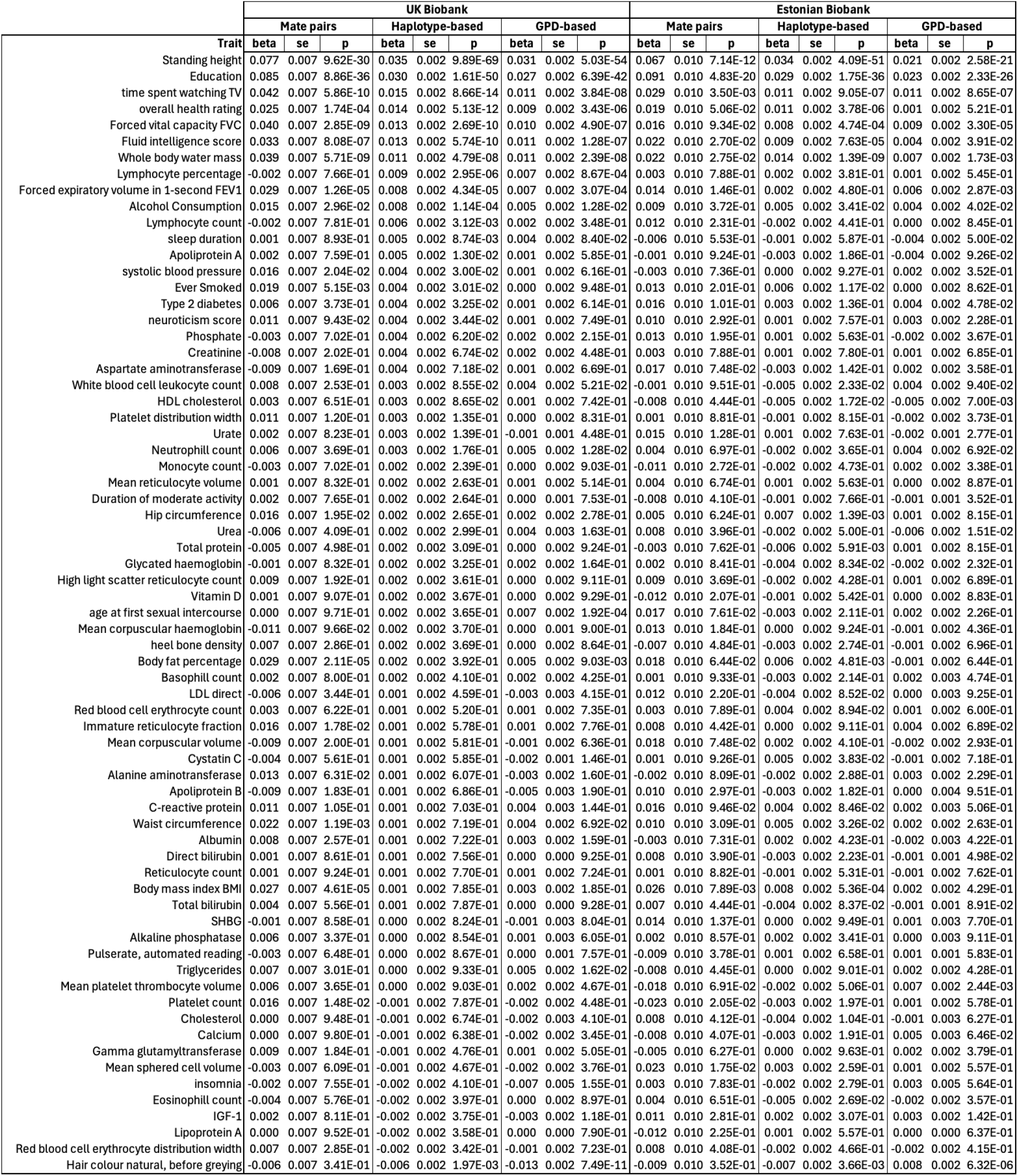
Genetic Assortative Mating in the UK and Estonian Biobanks. beta, effect size; se, standard error; p, P value.

